# The structured hairpin region of the bacterial ESCRT-III protein IM30 orchestrates stress-induced condensate formation

**DOI:** 10.64898/2026.02.28.708693

**Authors:** Ndjali Quarta, Katrin Debrich, Nadja Hellmann, Xingwu Ge, Pablo G. Argudo, Tika Ram Bhandari, Mischa Bonn, Martin Girard, Sapun H. Parekh, Lu-Ning Liu, Dirk Schneider

## Abstract

Biomolecular condensates are well characterized in eukaryotes, but their role in bacteria remains largely elusive. In the cyanobacteriumn *Synechocystis* sp. PCC 6803, the protein IM30, a member of the ESCRT-III superfamily of membrane remodeling proteins, forms stress-induced *puncta* across diverse environmental challenges, indicating a general adaptive response. Live-cell imaging reveals that IM30 is uniformly distributed throughout the cytoplasm at low concentrations but assembles into *puncta* upon exceeding a critical saturation threshold, a hallmark of liquid-liquid phase separation. Crucially, stress triggers *puncta* formation even below this threshold, suggesting stress lowers the phase-separation barrier. Super-resolution microscopy confirms spherical, condensate-like morphologies, while FRAP demonstrates rapid fluorescence recovery, consistent with with a fluid interior and dynamic exchange between *puncta* and the cytosol. Cellular IM30 levels exceed the *in vitro* determined critical concentration, placing the protein in a supersaturated state primed for condensation. Domain mapping identifies the structured α1–3 helical hairpin as the minimal phase separation driver; in contrast, the disordered α4–6 segment alone cannot phase-separate. Phase separation occurs within a physiologically relevant pH range (4.5–6.5), matching the localized acidification of the cyanobacterial cytoplasm at damaged thylakoid membranes. This directly links membrane stress, pH changes, and IM30 recruitment. Collectively, these findings establish IM30 *puncta* as *bona fide*, stress-responsive biomolecular condensates that function as rapid stress sensors and effectors, providing a mechanistic framework for phase separation and condensate formation by bacterial ESCRT-III proteins during environmental adaptation.

## Introduction

In eukaryotic cells, biochemical reactions are frequently organized and regulated by compartmentalization within membrane-bound organelles. As an alternative, cells can also form membraneless biomolecular condensates via liquid-liquid phase separation (LLPS) (Banani et al., 2017; Hyman et al., 2014). Within these dynamic, typically droplet-like structures, particular proteins and nucleic acids become temporarily concentrated, which enables precise spatial and temporal control of cellular activities. For example, under stress conditions, stress granules assemble to sequester mRNAs and halt their translation (Hofmann et al., 2021), whereas processing bodies (P-bodies) arise to sort and degrade transcripts (Brangwynne et al., 2009; Elbaum-Garfinkle et al., 2015). The formation of such condensates is typically triggered by specific cellular cues (Banani et al., 2017); nutrient deprivation, for instance, generates an excess of untranslated mRNA that drives P-body assembly.

The enrichment of specific biomolecules within a confined region while excluding others, a hallmark of LLPS, enables precise modulation of biochemical processes. For enzymes, this local concentration of reaction partners raises reaction rates in accordance with the law of mass action. In addition, the higher viscosity inside a condensate can influence catalytic speed, especially when the condensate’s visco-elastic properties change over time, for example, during maturation or solidification. Thus, LLPS provides a means to fine-tune enzymatic activity (Lyon et al., 2021; O’Flynn and Mittag, 2021). Likewise, in protein-protein interaction networks, such as cell-signaling pathways, concentrating key molecules in a condensate increases the probability of productive encounters, thereby shaping the signaling outcome (Su et al., 2021).

Condensates typically form when weak, multivalent interactions between proteins and/or nucleic acids become sufficiently strong to overcome the drive for molecular dispersion. This causes a homogeneous solution to separate into two coexisting phases, a dense, biomolecule-rich condensate and a surrounding dilute phase (Hyman et al., 2014).

Multivalent interactions between biomolecules permit the formation of a percolating network through thousands of transient contacts, a regime that is especially efficient in intrinsically disordered proteins (IDPs) or proteins containing intrinsically disordered regions (IDRs) (Borcherds et al., 2021). The lack of a rigid tertiary structure enables rapid re-organization of interacting regions, facilitating cooperative recruitment of binding partners. Nevertheless, ordered protein domains, such as SH3 or PDZ domains, can also drive LLPS when the interacting domains provide comparable multivalency (Cermakova and Hodges, 2023; Harmon et al., 2017; Hess and Joseph, 2025; Zeng et al., 2016; Zeng et al., 2018). Because condensate formation requires cooperative networking of a large number of molecules, LLPS exhibits a critical saturation concentration threshold (c_sat_) (Mittag and Pappu, 2022). This depends on the net interaction strength, which is modulated by a suite of biochemical parameters. Increases in protein abundance raise the valence density, while post-translational modifications such as phosphorylation, methylation, acetylation, or ubiquitination may alter a biomoleculés charge and polarity, thereby shifting the interaction parameter. Environmental variables, including pH, temperature, and ionic strength, affect electrostatic screening and hydrophobicity, moving the system within the co-existence phase space. Additionally, co-solutes, such as metabolites or crowding agents, can act as “co-condensers” lowering the saturation concentration (c_sat_) by providing extra interactions or by modulating solvent quality (Brzezinski et al., 2024).

The material properties of condensates are not always static. Initially, rapid exchange of constituents with the surrounding medium is possible in condensates, yet over time, the network might mature, resulting in a viscoelastic gel-like state. In extreme cases, irreversible structural rearrangements can give rise to solid-like, amyloid, or fibrillar aggregates (Alberti and Hyman, 2021; Galvanetto et al., 2025). Importantly, external cues can accelerate gelation, thereby sequestering specific enzymes or RNAs, whereas restoration of homeostatic conditions can dissolve the condensates by reversing the underlying interactions. This pronounced sensitivity to the cellular environment reflects a delicate balance between cohesive forces that drive assembly and forces that favor dispersion. Stress-related alterations in macromolecular concentration, temperature, pH, or ionic strength may tip this balance, prompting the swift appearance or disappearance of condensates and thereby enabling precise spatiotemporal regulation of metabolic and signaling pathways.

While initially thought to be an exclusive feature of complex eukaryotes, condensates are now increasingly recognized to occur across a broad spectrum of life, including fungi, protists, and even bacteria (Azaldegui et al., 2021; Staples et al., 2023; Stevens and Lasker, 2025). This widespread presence strongly suggests that organizing the cytoplasm without membranes via condensate formation is not a recent evolutionary innovation, but rather an ancient and fundamental mechanism. Bacterial systems illustrate the deep conservation of the general principles guiding condensate formation (Stevens and Lasker, 2025). For example, bacterial RNP (BR)-bodies share many properties with eukaryotic P-bodies and stress granules. BR-bodies are scaffolded by defined proteins such as RNase E, a principal mRNA-decay nuclease, and they incorporate additional proteins and RNAs required for efficient mRNA turnover (Al-Husini et al., 2020; Muthunayake et al., 2020). Bacteria likewise employ condensates for genome protection. For instance, *E. coli* induces the DNA-binding protein Dps to compact chromosomal DNA during the stationary phase (Gupta et al., 2023; Janissen et al., 2018), while *Pseudomonas aeruginosa* assembles polyphosphate condensates along its chromosomes under nitrogen starvation (Henry and Crosson, 2013; Racki et al., 2017).

Beyond the above-mentioned nucleic acid management, microorganisms exploit condensates for proteostasis, metabolism, and signaling. Quality-control condensates preserve proteome integrity, for example, by gathering misfolded proteins into polar aggresomes under stress (Jin et al., 2021). Metabolic condensates can improve pathway efficiencies, as illustrated by the RuBisCO-rich carboxysomes of cyanobacteria, which concentrate CO₂ for photosynthesis (Wang et al., 2019). Signaling condensates create spatially restricted hubs. For example, *Caulobacter crescentus* uses PopZ condensates to orchestrate opposing cell-fate determinants at each pole (Lasker et al., 2020).

While recent research confirms that condensates constitute a universal organization strategy across pro- and eukaryotes, bacterial condensates remain far less explored than their eukaryotic counterparts, and current evidence suggests that bacteria employ a comparatively limited molecular toolkit to generate multifunctional condensates, even though the underlying regulatory principles appear to be deeply conserved in pro- and eukaryotes.

The inner membrane-associated 30 kDa protein (IM30), also known as vesicle-inducing protein in plastids 1 (Vipp1), exemplifies how membrane remodeling proteins may interface with condensate biology. IM30 is essential for thylakoid membrane (TM) dynamics in cyanobacteria and chloroplasts (Fuhrmann et al., 2009b; Kroll et al., 2001; Westphal et al., 2001; Zhang et al., 2012). IM30 is dispersed throughout the bacterial cytoplasm yet forms dynamic *puncta* near internal membranes under light stress (Bryan et al., 2014; Gates et al., 2022; Gutu et al., 2018), similar to the stress-induced formation of *puncta* at the cytoplasmic membrane observed for other bacterial homologues (Dominguez-Escobar et al., 2014; Engl et al., 2009; Yamaguchi et al., 2013). Although the precise nature of these assemblies remains uncertain, the proteins seem to localize near internal membranes, particularly TMs, consistent with a potential role in membrane stabilization and/or repair (Bryan et al., 2014; Gates et al., 2022; Gutu et al., 2018; Junglas and Schneider, 2018). Isolated IM30 has been shown to undergo phase separation and form condensates *in vitro* under physiological conditions of protein concentration and ionic strength, potentially linking the *puncta* observed *in vivo* to biomolecular condensates (Quarta et al., 2026). *In vitro*, condensate formation of IM30 is largely driven by electrostatic interactions, exhibiting the typical re-entrant phase behavior of polyampholytes. Isolated IM30 monomers have seven predicted α-helical regions (helices α0-α6) (Gupta et al., 2021; Junglas et al., 2025; Liu et al., 2021; Schlösser et al., 2023) (Figure 1A), albeit when free in solution, monomeric IM30 appears to form only three stable α-helices, α1-α3, with α3 being a direct extension of α2 (Heidrich et al., 2016; Junglas et al., 2020; Quarta et al., 2024). The helices α1 and α2/3 interact to form a stable helical hairpin structure. The remaining regions are unstructured in the IM30 monomeric state, *i.e.* ∼ 50% of IM30 monomers constitute an IDR (Junglas et al., 2020; Quarta et al., 2024). *In vitro*, at least, isolated IM30 monomers spontaneously oligomerize in solution to form diverse barrel or rod structures (Fuhrmann et al., 2009a; Gupta et al., 2021; Junglas et al., 2025; Liu et al., 2021; Siebenaller et al., 2021) (Figure 1B). This structure formation is driven by the multiple interactions IM30 monomers can establish between individual helices in higher oligomeric barrel structures, where a single monomer can interact with up to 16 different monomers in the same barrel (Gupta et al., 2021). Helices form an interwoven network with conserved intermolecular interfaces (Schlosser et al., 2023). Thus, multivalency, a hallmark of condensate formation, is inherent in IM30 proteins. Upon membrane contact, IM30 forms large polymeric structures *in vitro*: carpets, barrels, rods or spirals (Fuhrmann et al., 2009a; Gupta et al., 2021; Junglas et al., 2022; Junglas et al., 2025; Junglas et al., 2020; Liu et al., 2021; Naskar et al., 2025; Pan et al., 2024; Schlösser et al., 2025). Membrane-bound barrels and rods can enclose and thereby tubulate the membrane, which creates high curvature, and this likely accounts for the strong propensity of IM30-covered membranes to fuse spontaneously, a feature IM30 shares with ESCRT-III superfamily proteins (Hennig et al., 2015; Junglas et al., 2025). In fact, recent structural analyses revealed that IM30 and homologous bacterial proteins are members of an ESCRT-III superfamily, previously thought to be exclusive to eukaryotes (Gupta et al., 2021; Junglas et al., 2021; Liu et al., 2021; Schlösser et al., 2023). In eukaryotes, the endosomal sorting complex required for transport (ESCRT) plays pivotal roles in diverse membrane remodeling processes, including vesicle formation, cytokinesis, and membrane repair (Henne et al., 2011; Remec Pavlin and Hurley, 2020; Vietri et al., 2020). In these processes, the ESCRT-III polymers trigger membrane budding and scission through dynamic polymer formation, reorganization, and disassembly (Pfitzner et al., 2021). All ESCRT-III superfamily members share a conserved core of five α-helical regions (α1–α5), with the α1–3 hairpin forming the structural scaffold, a feature also observed in IM30 (McCullough et al., 2018; Schlösser et al., 2023). Besides the conserved helices α1-5, several ESCRT-III proteins contain an additional N-terminal amphipathic helix (α0) crucial for membrane binding (Buchkovich et al., 2013; Gupta et al., 2021; Huber et al., 2020; Hudina et al., 2025; Junglas et al., 2025; Liu et al., 2021; Schlösser et al., 2025), and eukaryotic ESCRT-IIÍs may contain additionally C-terminal regions, many of which are involved in protein-protein interactions (Banjade et al., 2021; Nguyen et al., 2020; Obita et al., 2007; Shim et al., 2007; Tan et al., 2015).

**Figure 1.**
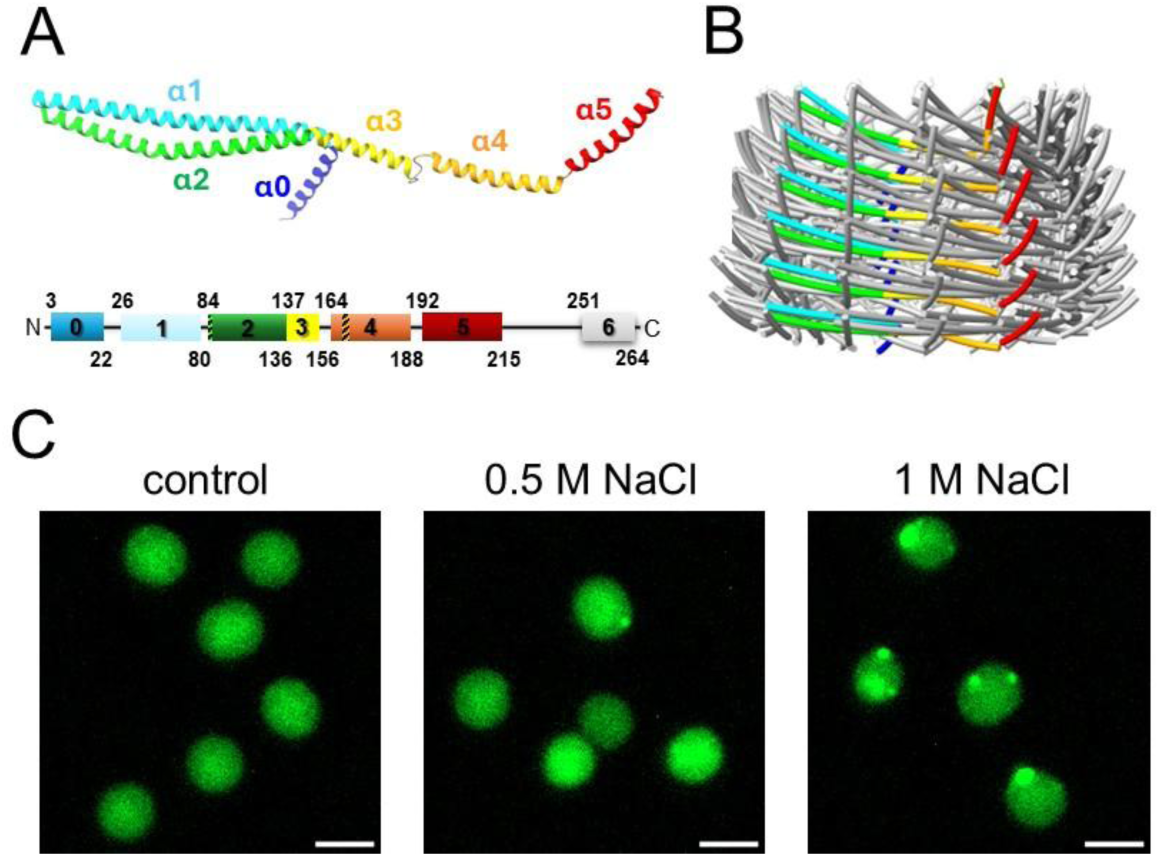
IM30 forms *puncta* structures in salt-stressed *Synechocystis* cells. (A) Structure of a wild type (wt) IM30 monomer. Top: monomeric IM30 extracted from an oligomeric assembly (PDB 703Y); helix α6 is absent from the resolved model. Bottom: domain illustration of the wt protein with helix numbering. The numbers indicate the start and end-residue positions used to express truncated IM30 variants. Regions mutated in the IM30* construct are marked by a dashed bar. (B) Representative C16-symmetric IM30 barrel (PDB 703Y). (C) Fluorescence microscopy of *Synechocystis* cells expressing mVenus-tagged IM30. Images show cells before (control) and after the addition of 0.5 M or 1 M NaCl. Salt stress induces the formation of *puncta* (overview images of multiple cells are provided in Figure S1). Scalebar. 2 µm.

Eukaryotic ESCRT-IIIs and IM30 interact with ATPases (Vps4 and Hsp70/DnaK, respectively) (Babst et al., 1998; Bryan et al., 2014; Liu et al., 2007; Maity et al., 2019), triggering oligomer disassembly, which is in IM30 coupled with substantial monomer unfolding and exposure of IDRs at both termini (α0 and α4-6) while preserving the α1-3 structural core (Quarta et al., 2024).

To summarize, IM30 monomers are multivalent, roughly 50% disordered, may experience further unfolding based on interactions, and undergo phase separation at physiological protein and salt concentrations *in vitro* (Quarta et al., 2026). This suggests that the condensates observed *in vitro* and *puncta* structures that appear in living cyanobacterial cells under light- or salt-stress may be connected though the nature of the *puncta* remains elusive.

Here, we demonstrate that the cyanobacterial ESCRT-III superfamily member IM30 forms discrete *puncta* in living cells that exhibit defining features of biomolecular condensates. *In vitro*, purified IM30 undergoes LLPS upon mild acidification, forming spherical droplets with a *c*_sat_ that is well below the estimated intracellular concentration in living *Synechocystis* cells, making spontaneous condensation thermodynamically favorable. *In vivo*, IM30 is diffusely distributed in the cytoplasm under basal conditions. However, when cellular protein levels rise, or in response to stressors such as low pH, elevated NaCl, or heat, IM30 rapidly assembles into spherical *puncta*. Fluorescence recovery after photobleaching (FRAP) of a IM30–mVenus fusion expressed in living bacterial cells revealed rapid fluorescence recovery, confirming a dynamic, fluid interior and continuous exchange of monomers with the surrounding cytosol. Mutational analysis identified the α1–3 helical hairpin, a conserved structural core across all ESCRT-III family members, as both necessary and sufficient for phase separation. Truncated variants lacking this hairpin fail to condense, whereas an oligomerization-deficient IM30 variant still forms *puncta*, proving that barrel assembly is dispensable for condensation.

Physiologically, moderately lowering the pH triggers *in vitro* and *in vivo* phase separation, which matches the locally acidified microenvironment created at damaged TMs, where proton leakage lowers pH spatiotemporally. Thus, membrane stress can locally induce LLPS, providing a rapid, reversible reservoir of monomeric IM30.

## Results

### In vivo formation of IM30 puncta from IM30 monomers

In the absence of stress, C-terminally mVenus-labeled wt IM30 is uniformly distributed throughout the cyanobacterial cytoplasm (Figure 1C). However, upon exposure to 0.5 M or 1 M NaCl, the protein assembles into discrete *puncta*. At 0.5 M NaCl, approximately 15% of cells exhibit such structures, typically one *punctum* per cell. At 1 M NaCl, over 60% of cells contain *puncta*, with roughly 50% cells containing a single *punctum* and about 15% containing two *puncta*. These structures emerge immediately upon adding *Synechocystis* cells to high-salt medium (Figure 1C).

Previously, IM30 phase separation has been observed *in vitro* at elevated salt concentrations, yet solely in the presence of a molecular crowder to induce condensate formation (Quarta et al., 2026). In contrast, within a cellular environment, elevated salt concentrations alone appear sufficient to drive phase separation.

Despite their rapid formation, the precise nature of the forming *puncta* remains unclear. Nevertheless, the intrinsic ability of monomeric IM30 to undergo phase separation and form condensates under physiological protein and salt concentrations (Quarta et al., 2026) strongly suggests that these *puncta* are biomolecular condensates formed by LLPS.

Key characteristics have previously been defined that allow considering such assembly structures as a result of phase separation (Mittag and Pappu, 2022), including the formation of condensates upon reaching a saturation concentration (c_sat_) and a difference in component density between two phases. To address these challenges, a robust experimental framework has recently been established to identify, assess, and characterize biomolecular condensates in living bacteria (Hoang et al., 2024). Induced (over)expression of a fluorescently labelled protein is a simple and quick method for gaining first insights into the material state of the IM30 *puncta* forming in living bacterial cells.

To investigate the *in vivo* behavior of fluorescently labeled IM30 in living cells, we first expressed the IM30 wt protein C-terminally fused to mVenus in *E. coli*, and monitored accumulation of mVenus fluorescence in the cytoplasm (Figure 2A, B). Although no mVenus fluorescence was initially detectable (Figure 2B, 0h), within just 1 h of inducing gene expression, fluorescence appeared throughout the bacterial cytoplasm, accompanied by the spontaneous formation of *puncta* structures (Figure 2B, 1h). One hour post-induction, essentially all (97%) IM30-mVenus-expressing cells displayed *puncta*, with an average of 1–2 *puncta* per cell. Yet, although no significant background fluorescence was apparent under standard imaging conditions before the addition of IPTG, increasing the contrast revealed a uniform, low-level mVenus signal distributed throughout the entire cytoplasm of all cells, already in uninduced cells (Figure S2). This indicates that the concentration of IM30–mVenus had already surpassed a concentration sufficient to drive condensate formation shortly after induction of protein expression. Thus, two IM30 phases appear to co-coexist in living bacteria, a hallmark of phase separation. Notably, quantifying the ratio of free (cytoplasmic, dilute-phase) IM30–mVenus to that sequestered in *puncta* (condensate phase), *i.e*., estimating an *in vivo* saturation concentration (*c*_sat_), was not feasible, as the *puncta* exhibited fluorescence intensities too high to allow reliable background subtraction or intensity calibration.

**Figure 2:**
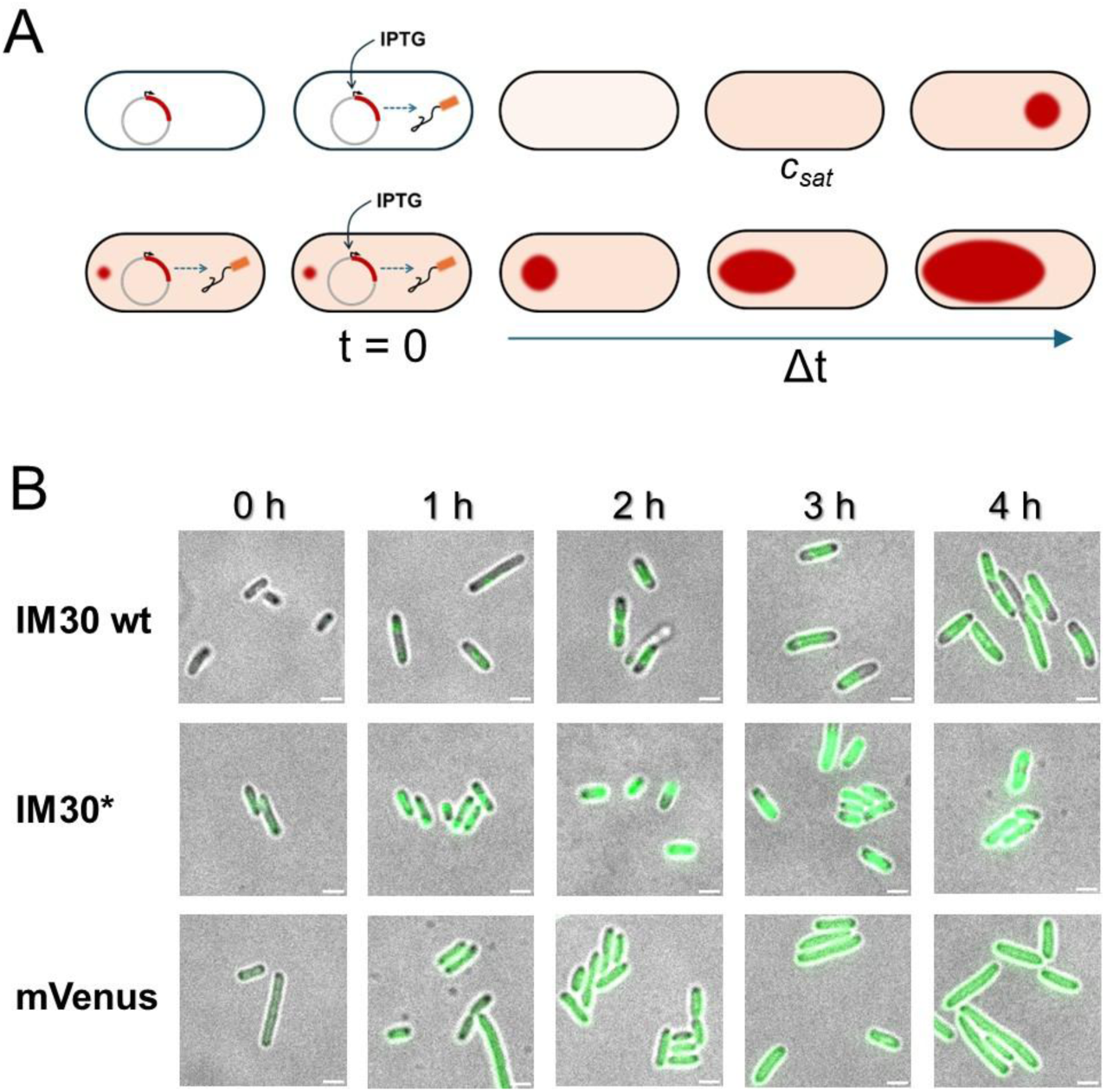
Formation of IM30 *puncta* in *E. coli*. (A) IM30(*) and free mVenus were expressed in *E. coli* under inducible control. Upper panel: When basal expression levels are low, and c_sat_ is sufficiently high, *puncta* form only after sufficient protein has accumulated in the cytoplasm following induction. Lower panel: When basal expression is high and c_sat_ is low, *puncta* may appear even in uninduced cells, reflecting spontaneous condensate formation already at baseline expression levels. (B) mVenus-tagged IM30 wt, IM30*, and free mVenus were expressed in *E. coli* under control of the T7 promoter. For free mVenus and IM30*, the basal expression was high. mVenus distributed diffusely throughout the cytoplasm, while IM30* formed *puncta* even without induction. For IM30 wt, basal expression was lower, and small *puncta* became visible 1 h after induction of gene expression. Merged pictures of phase contrast and mVenus fluorescence are shown. Non-merged overview images showing multiple cells are provided in Figure S3. Scale bar: 2 µm.

Upon induction of gene expression, IM30 wt *puncta* formed rapidly, grew in size over time, and ultimately occupied a substantial portion of the bacterial cytoplasm (Figure 2B).

Similarly, when free mVenus was expressed under identical conditions, a low cytoplasmic fluorescence was observed in non-induced cells, yet the fluorescence increased gradually following induction of gene expression. However, no *puncta* were observed. This confirms that *puncta* formation is specifically mediated by IM30 (Figure 2B).

IM30*, an IM30 variant where four amino acids were replaced in helix α4 that establishes a crucial contact between IM30 monomers (Figure 1A), does not form large, ordered protein supercomplexes anymore (Heidrich et al., 2016; Junglas et al., 2020). Importantly, the helical propensities of the individual regions are not altered (Quarta et al., 2024), making IM30* an excellent model for monomeric IM30. We previously demonstrated that IM30**\*** still undergoes phase separation, at least under the *in vitro* conditions (Quarta et al., 2026). When expressed in *E. coli*, *puncta* formation within the cytoplasm was observed already prior to induction of gene expression (Figure 2B, 0h). Upon induction of expression, *puncta* started to grow comparable to the *puncta* formed by IM30 wt.

As cellular metabolism and protein turnover are much slower in *Synechocystis* than in *E. coli*, and *Synechocystis* cells have a high background fluorescence, making time-lapse experiments less feasible. Therefore, we used another strategy to determine whether the formation of IM30 condensates is also a response to cellular protein concentration in living cyanobacterial cells. An IM30-mVenus fusion protein was ectopically expressed in living cells, and gene expression was controlled by a rhamnose-inducible promoter. Twenty-four hours after induction of gene expression by adding increasing concentrations of rhamnose, *Synechocystis* cells were analyzed by fluorescence microscopy. As shown in Figures 3A and S4, the intracellular fluorescence intensity increased with increasing rhamnose concentrations in cells expressing IM30–mVenus, yet even at the highest inducer level, no *puncta* were observed. However, when these same cells, which exhibited uniform cytoplasmic fluorescence, were subjected to salt stress, *puncta* readily formed, as previously demonstrated (Figure 1C), even at low inductor concentrations. This indicates that high-level expression of IM30-mVenus alone is insufficient to drive spontaneous *puncta* formation in living *Synechocystis* cells, and *puncta* formation requires an additional trigger, here salt stress. Unfortunately, the high background fluorescence of the cyanobacterial cells hindered the determination of a meaningful partitioning ratio and c_sat_.

**Figure 3:**
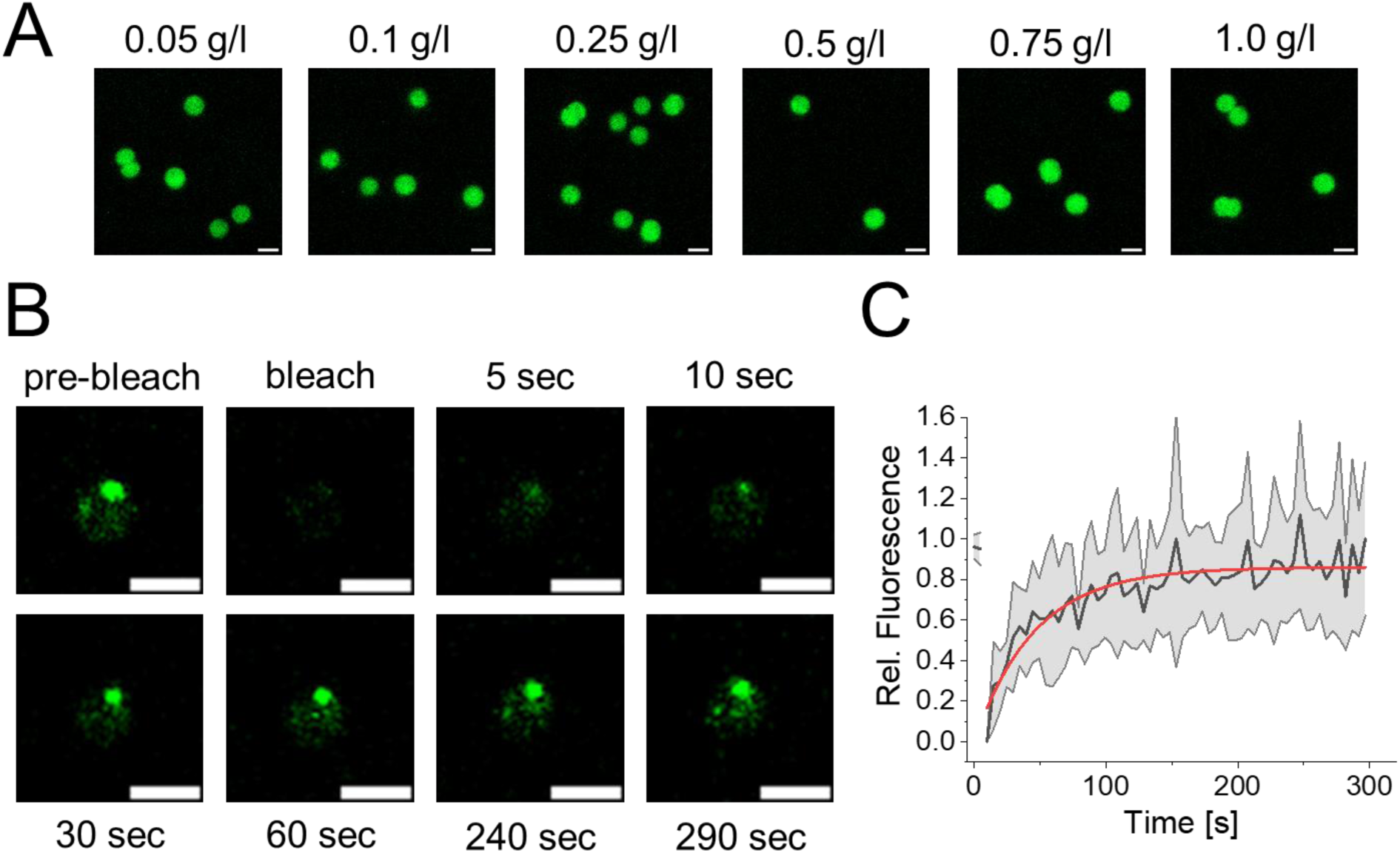
Formation and characteristics of IM30 *puncta* in *Synechocystis*. (A) Fluorescence micrographs of *Synechocystis* cells expressing IM30-mVenius upon addition of increasing concentrations of the inducer rhamnose. Scale bar: 2 µm. Due to the high background fluorescence, quantification of the increase in intracellular mVenus signal was not straightforward. Quantification of intracellular fluorescence is presented in Figure S4. (B, C) FRAP of IM30-mVenus. (B) Fluorescence microscopy images of a fully bleached salt-induced IM30 condensate in *Synechocystis* before and after the bleach pulse. Non-bleached condensates in other cells were used as controls to correct for acquisition bleaching. The scale bar in (A) and (B) is 2 µm. (C) Normalized fluorescence intensity measured in FRAP experiments with *in vivo* IM30 condensates. Error bars represent SD (n=3 biological replicates (N = 6 cells)).

Collectively, our observations in living bacterial cells, *E. coli* or *Synechocystis*, clearly suggest that IM30 condensates form in response to increasing intracellular IM30 concentrations and salt stress. Importantly, these condensates coexist with a dilute cytoplasmic phase, a hallmark feature of phase-separated biomolecular condensates.

To probe the exchange between condensates and the cytosol, we turned to live-cell imaging of IM30 *puncta* in *Synechocystis*, tracking their dynamics via the fluorescence recovery after photobleaching (FRAP) of individual condensates. *Puncta* formation was induced via the addition of 1 M NaCl to cells expressing IM30-mVenus (see Figure 1C). In the FRAP experiments (Figure 3B, C), the *puncta* fluorescence showed an almost complete recovery after 3 min, and the condensates were able to recover approximately 90% of their initial fluorescence signal via a dynamic exchange of the (bleached) IM30-mVenus proteins with the surrounding cytoplasm (Figure 3B, C). From the recovery curves (Figure 3C) we were able to calculate the time it took for the fluorescence intensity to recover to half of its final value, *i.e.* the recovery half-time (*t_1/2_*), which was approximately 37.1 ± 20.0 s, as well as a diffusion coefficient *D* of 0.0033 +/- 0.0017 µm^2^/s. These exchange dynamics are characteristic of biomolecular condensates formed *in vivo* via LLPS, confirming their dynamic nature, rather than stable protein aggregation.

As IM30 spontaneously forms large supercomplexes which dynamically form and disassemble, at least in solution (Carlton and Baum, 2023; Christ et al., 2017; Pfitzner et al., 2021; Schlosser et al., 2023), the exchange dynamics observed in living cells could simply signify exchange of protomers in large polymeric structure, barrels or rods, or super-assemblies of such complexes. Thus, to test whether the formation of higher-order oligomers is required for the *in vivo* formation of *puncta*, we ectopically expressed C-terminally mVenus-labeled IM30* (IM30*-mVenus) in *Synechocystis*. When analyzing living *Synechocystis* cells using super-resolution fluorescence microscopy, upon expression of IM30 wt as well as mVenus-labeled IM30* we observed the formation of fluorescent *puncta* (Figure 4A) in living *Synechocystis* cells, indicating that the disrupted contacts responsible for forming and stabilizing IM30 barrels and rods are not vital for *puncta* formation *in vivo*, in line with the observations made in *E. coli* (Figure 2). While *puncta* formation appears to be less pronounced in the case of IM30* when compared to the wt (Figure 4A, B), the observations demonstrate that *puncta* formation does not rely on IM30’s propensity to form large, structured supercomplexes. Notably, some IM30* appears to bind to intracellular membranes, indicated by a halo lining the cytoplasmic membrane, which is consistent with recent work showing that the α1-3 helical hairpin, that is exposed in IM30*, binds to membrane surfaces (Schlösser et al., 2025). Furthermore, binding of IM30* monomers to internal membranes also reduces the number of free oligomers and monomers in solution, which presumably affects *puncta* formation *via* changing the concentration of available IM30* (monomers) in the cytoplasm.

**Figure 4:**
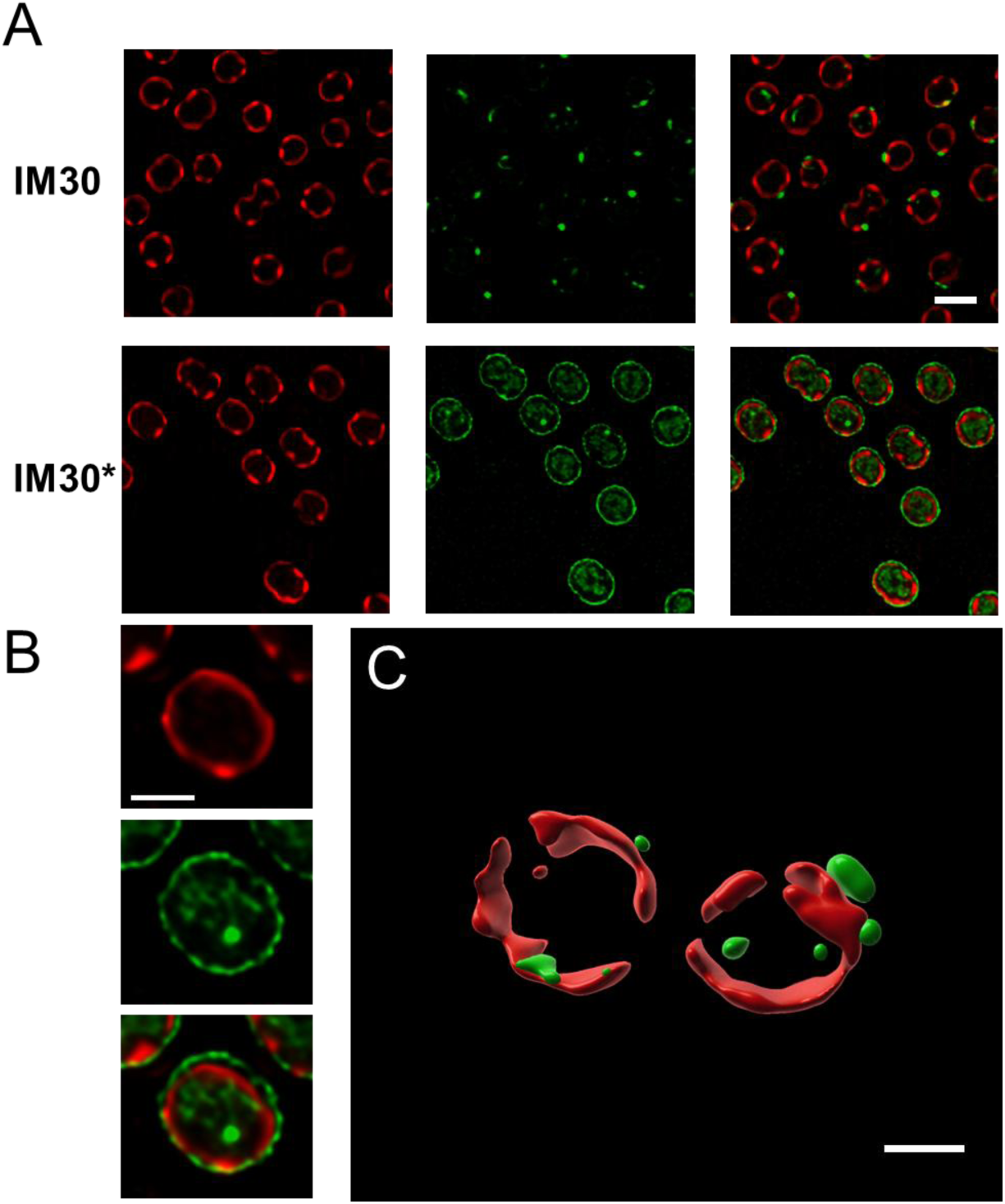
IM30 forms spherical structures in living cells. (A) Super-resolution fluorescence images showing *Synechocystis* cells expressing mVenus-tagged IM30 wt and IM30*. Chlorophyll signals representing TMs are colored in red (1^st^ column) and mVenus signals colored green (2^nd^ column). Scale bar = 2 μm. (B) Zoom-in image of a *Synechocystis* cell expressing mVenus-tagged IM30*. An IM30 *punctum* as well as protein attached to the cytoplasmic membrane are visible. Scale bar = 1 μm. (C) 3D-rendering of TM (red) and mVenus (green) fluorescence signals show that IM30 proteins form spherical assemblies. Scale bar = 1 μm.

Fluorescence image stacks and 3D-rendered images of *Synechocystis* cells expressing IM30-mVenus revealed that the *puncta* forming *in vivo* were situated near TMs (Figure 4C); however, they mostly do not appear to represent membrane-coating carpets, at least under the tested experimental conditions. 3D reconstruction of the condensate from fluorescence Z-stacks (0.11 μm spacing) revealed an apparently spherical body, resembling biomolecular condensates formed *via* protein phase separation. As with the wt protein, IM30* also formed spherical structures *in vivo*, in living cyanobacterial cells, revealing that the contacts responsible for forming and stabilizing IM30 wt barrels and rods are not vital for the *in vivo* formation of spherical IM30 structures.

Together, IM30 assembles into spherical structures in living cyanobacterial cells. These structures form either when intracellular IM30 concentration exceeds its critical saturation threshold or when environmental cues, in this case elevated salt, shift the phase behavior of the protein. Within the cell, IM30 partitions between a soluble, dispersed cytoplasmic pool and a condensed phase, with monomers exchanging between the two compartments, a hallmark of dynamic, liquid-like condensates forming due to LLPS. Importantly, the interactions driving the formation of these spherical condensates likely differ from those that stabilize the highly ordered, polymeric IM30 structures (*e.g.*, rings or barrels) observed under other conditions.

### The structured helical hairpin region orchestrates dynamic self-assembly and condensate formation

IM30 forms liquid condensates *in vitro* at physiologically relevant protein and salt concentrations (Quarta et al., 2026) as well as *in vivo* when protein concentrations are >c_sat_ and/or upon (at least) salt stress (Figures 1-3). However, it remains unknown which part of the protein mainly mediates LLPS.

IDRs often possess inherent flexibility that facilitates multiple transient interactions, leading to LLPS and condensate formation (Pappu et al., 2023). While the C-terminal IM30 part, with helices α4-6, as well as helix α0 are indeed disordered in monomeric IM30, the central helical hairpin formed by helices α1-3, remains stably folded (Junglas et al., 2020; Quarta et al., 2024). To pinpoint the region(s) mainly driving IM30 phase separation, we next monitored and compared *in vitro* condensate formation of monomeric IM30* and truncated IM30 variants (Figure 5A, B).

**Figure 5:**
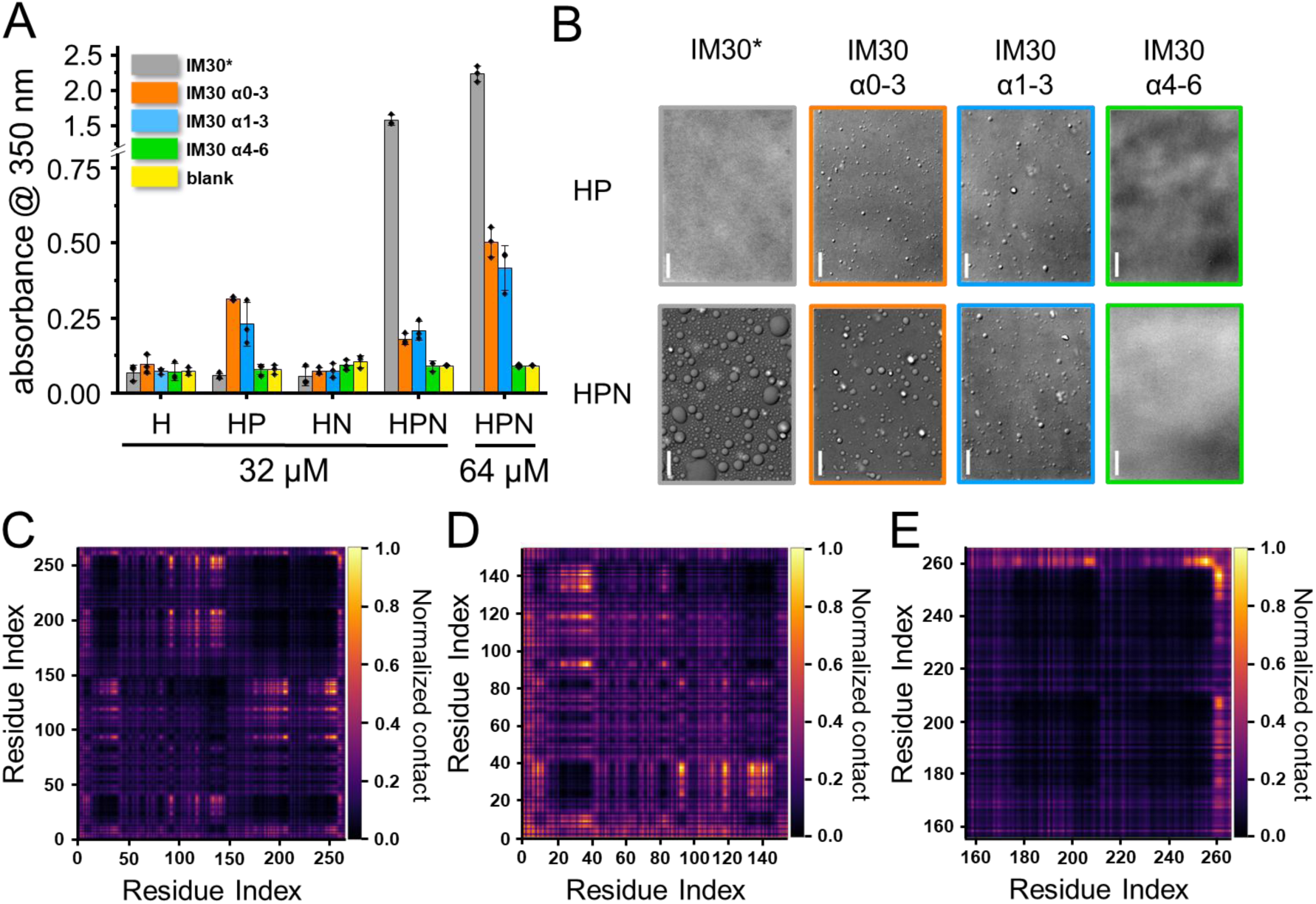
Condensate formation of IM30 variants. (A) Turbidity of solutions containing 32 µM or 64 µM IM30* (grey), IM30_α0-3_ (orange), IM30_α1-3_ (blue), IM30_α4-6_ (green), or no protein (yellow). Formation of condensates was tested in 20 mM HEPES (pH 7.5) without additives (H), with 10% PEG (HP), 100 mM NaCl (HN), or both (HPN) (n=3, SD). For the beginning and end of the indicated α-helices, see Figure 1A. (B) DIC microscopy images of protein solutions in buffer HP or HPN. Frame colors are as columns in (A). For the HP measurements, the protein concentration was 32 µM, for HPN 64 µM. Scale bars are 10 µm. An SDS-PAGE gel of the purified proteins is shown in Figure S5. (C-E): Interchain contact map of IM30 condensates at 220 K and κ = 0.097 for the full-length wt IM30 (C), α0-3 (D) and α4-6 (E). A contact map for full-length IM30* is shown in the supplement (Figure S6).

Recently, we have shown that IM30*, as well as urea-destabilized IM30 wt, forms condensates via LLPS at a physiological pH of 7.5 when 100 mM NaCl and 10% PEG were added (Quarta et al., 2026). As observed now, all (truncated) protein variants containing the stable α1-3 helical hairpin, *i.e.,* IM30*, IM30_α0-3_, and IM30_α1-3_, phase separated, with the condensate-forming propensity of IM30_α0-3_, and IM30_α1-3_ being less pronounced than in the case of full-length IM30* (Figure 5A, B). The droplet sizes observed via DIC imaging, as well as the turbidity values of the solutions containing the truncated proteins, were significantly lower than those of IM30* at equal protein concentrations (Figure 5A, B). As we did not observe any significant differences in turbidity or condensate structure between IM30_α1-3_ and IM30_α0-3_, helix α0 appears not to substantially affect condensate formation, at least not *in vitro*. Importantly, the isolated IDR IM30_α4-6_ did not form condensates under any of the tested conditions.

However, while IM30* did not phase separate in buffer containing solely PEG (Quarta et al., 2026) but no NaCl, the truncated proteins featuring the stable α1-3 helical hairpin region, *i.e.,* IM30_α0-3_ and IM30_α1-3_, formed condensates already when solely 10% PEG was added (Figure 5A). This suggests that NaCl is less crucial for the phase separation of these proteins compared to IM30*, which may be explained by the coacervation of polyampholytes (Chang et al., 2017; Das and Pappu, 2013; Sing and Perry, 2020). Coacervation is essentially linked to three polymer properties: the blockiness of the charge pattern (characterized by kappa) (Das and Pappu, 2013), the fraction of charged residues (FCR), and the net charge per residue (NCPR). All the sequences considered show similar charge patterns (kappa ∼ 0.15 – 0.18 Å^-1^), which is sufficiently large to collapse the chain (Das and Pappu, 2013). However, α1-3 and α0-3 are strong polyampholytes (FCR ∼ 0.33), whereas α4-6 is weak (FCR∼ 0.2), leading to stronger electrostatic interactions for the former. In addition, α4-6 carries a net charge (NCPR∼0.11), as opposed to the almost neutral α0-3 and α1-3 (∼0.03).

To gain further insights into which regions of the protein drive phase separation, we next calculated the frequency of inter-chain contacts between all residue pairs within the condensates using coarse-grained molecular dynamics, as illustrated in the contact maps (Figure 5C-E). The off-diagonal contact map of the full-length wt protein (Figure 5C) as well as of IM30* (Figure S6) reveals a substantial number of contacts between residues in the helical-hairpin α1-3 and the C-terminal α4-6 regions in condensates. Based on the experimental observations, we subsequently tested the individual α0-3 and α4-6 segments in our simulations. Figure 5D shows a high contact probability among residue pairs within the N-terminal α0-3 region, indicating extensive intermolecular interactions in this domain, in excellent agreement with the experimental results. In contrast, the C-terminal region (α4-6) displays very low contact probability (Figure 5E), with only sparse interactions, mainly involving a few terminal residues. This is evident from the largely blank regions in the contact map. These results support the experimental observation that the α1-3 helical hairpin region alone is sufficient to drive condensate formation, whereas the C-terminal region by itself has limited intrinsic condensation capability.

Taken together, our observations suggest that the α1-3 helical-hairpin region, yet not the extended IDR (IM30_α4-6_), promotes condensate formation *in vitro* via coacervation. However, as full-length IM30* has an increased propensity to phase separate in the presence of NaCl and PEG compared to any of the truncated IM30 variants (Figure 5A, B), the extended IDR clearly promotes phase separation in the context of the full-length protein, in line with the additional contacts observed in our simulations (Figures 5C, S6).

To next test whether the *in vitro* finding that the α1–3 helical hairpin drives condensate formation also holds true *in vivo*, we next generated *Synechocystis* cells expressing mVenus-tagged truncated IM30 variants. This enabled us to directly visualize and compare the phase separation behavior of full-length and truncated IM30 within living cells. As IM30 is essential, chimeric genes encoding the mVenus-tagged proteins were expressed ectopically. Upon expression of the truncated variant IM30_α0-3_-mVenus, *puncta* formation was observed (Figure 6A), in perfect agreement with our *in vitro* observations (Figure 5). When the 3D structure of a *Synechocystis* cell expressing IM30_α0-3_-mVenus was rendered, a mesh was observed in some cells instead of single spherical droplet-like structures (Figure 6A, B). This could form due to sterically constrained binding forces that align the truncated proteins in specific orientations, favoring filament or sheet formation rather than isotropic coalescence. In the absence of the C-terminal tail, the structured helical-hairpins potentially interact with each other in a more defined orientation, which favors the formation of linear or sheet-like assemblies rather than isotropic packing. Alternatively, other polymers, especially the negatively charged RNA and DNA, might bridge the positively charged α1-3 hairpins, and the formed condensate therefore inherits the polymer’s contour (here: a loose web) rather than forming a compact sphere. This has, *e.g.,* been observed for DNA-histone structures (Zhou et al., 2025).

**Figure 6:**
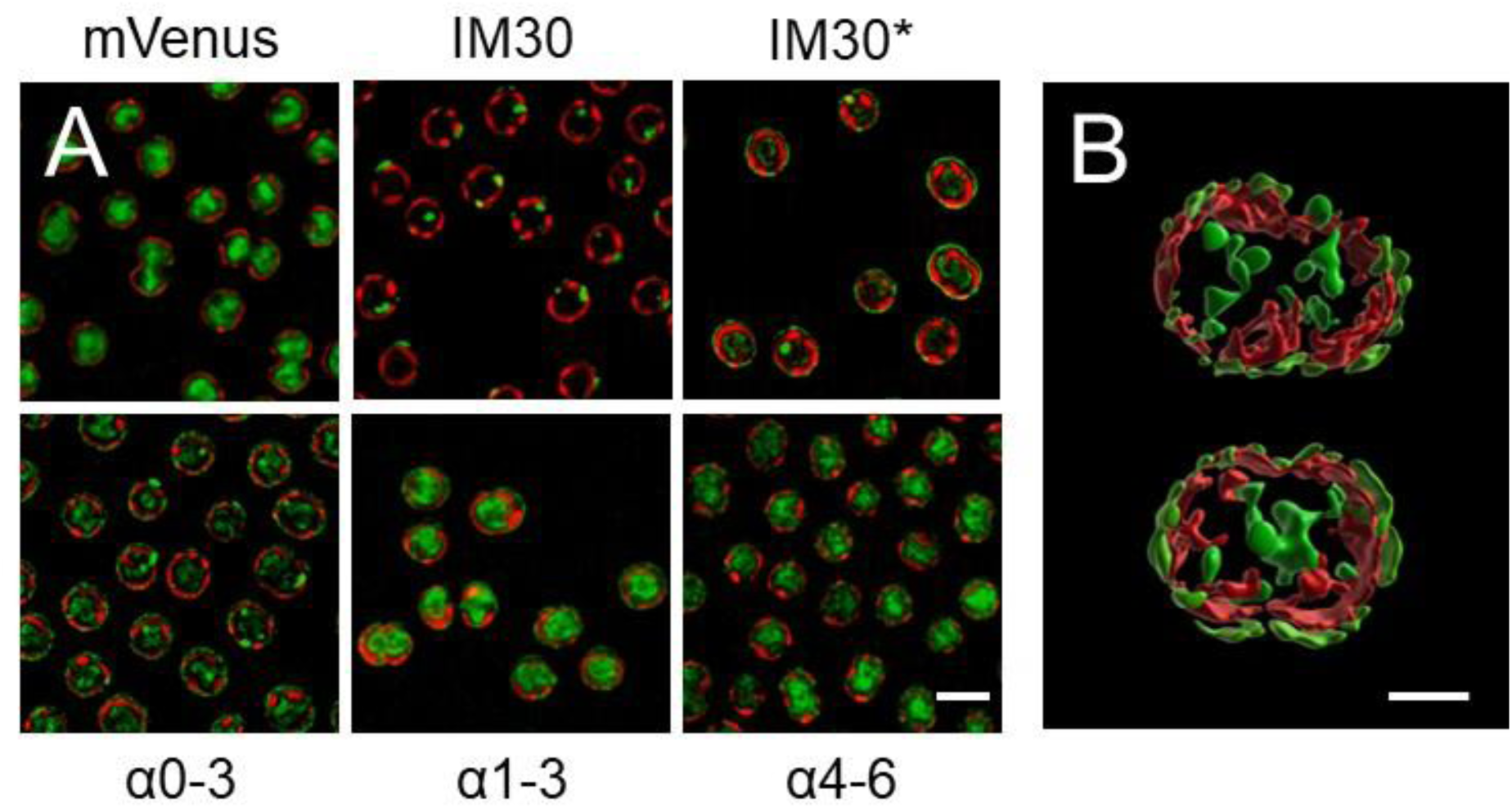
*Puncta* formation of IM30 variants in *Synechocystis* visualized via structured illumination microscopy. (A) Merged images showing TM fluorescence (red) as well as mVenus fluorescence (green, compare Figure 4) of *Synechocystis* cells expressing solely mVenus, IM30-mVenus, IM30*-mVenus, IM30_α0-3_-mVenus, IM30_α1-3_-mVenus, or IM30_α4-6_-mVenus. Scale bar: 2 µm. (B) 3D-rendered Synechocystis cell expressing mVenus fused to IM30α0-3. TM colored in red and mVenus signal in green. An increased formation of condensates is observed, which tend to fuse and form intracellular networks. Scale bar: 1 µm.

In contrast to IM30 wt, IM30*, or IM30_α0-3_, the fluorescence signals of IM30_α4-6_-mVenus or IM30_α1-3_-mVenus, respectively, were exclusively distributed in the cytosol (Figure 6A), suggesting that helix α0 is more important for *in vivo* condensate formation than *in vitro* (Figure 5). Together, our results revealed that the α1-3 helical-hairpin structure orchestrates condensate formation both *in vitro* and *in vivo*.

### The in vivo formation of IM30 puncta/condensates is a general stress response mechanism

Previously, IM30 *puncta* have been observed in living *Synechocystis* cells when cells were high-light- or salt-stressed (Bryan et al., 2014; Quarta et al., 2026). Similarly, the homologous bacterial proteins PspA and LiaH also form *puncta* structures in stressed cells (Dominguez-Escobar et al., 2014; Engl et al., 2009; Yamaguchi et al., 2013). The physiological functions of all these bacterial ESCRT-III proteins are linked to membrane repair, stabilization and/or dynamics, although the precise mechanism of action remains largely enigmatic. Since LLPS is increasingly recognized as a general stress-response mechanism for many proteins, we next asked whether IM30 *puncta* formation follows the same pattern, acting not only under salt or high-light stress, but also as a broader cellular response to diverse stressors.

Upon treatment of *Synechocystis* cells with diverse stressors, we observed the formation of *puncta* in most cases (Figures 7, S1 and S7). Except for lowering the temperature (from 30°C to 20°C) on cells, we observed a dramatic formation of *puncta* structures for all other stressors. We note that addition of 0.1 M maltose (to increase the solutiońs osmolarity) or slightly increasing the temperature to 40°C showed an attenuated increase the number of *puncta* observed in treated cells compared to the other stressors. Treating the cells with 5% ethanol, 0.25 mM H_2_O_2_, 0.5 M maltose, and 0.5 M NaCl moderately induced the formation of (typically) a single *punctum* in 10-20% of all cells analyzed. Notably, pronounced *puncta* formation was not limited to salt stress: exposure to 45°C (heat stress) also triggered robust *puncta* formation. Moreover, the frequency of *puncta* per cell increased, shifting from a single to multiple *puncta*, suggesting a dose-dependent response to stress intensity. As mentioned above, in cells stressed with 1 M NaCl, about 15% of all stressed cells contained two *puncta* and 2% even three. Together, these analyses clearly show that *puncta* formation appears to be a more general stress response mechanism in *Synechocystis*.

**Figure 7:**
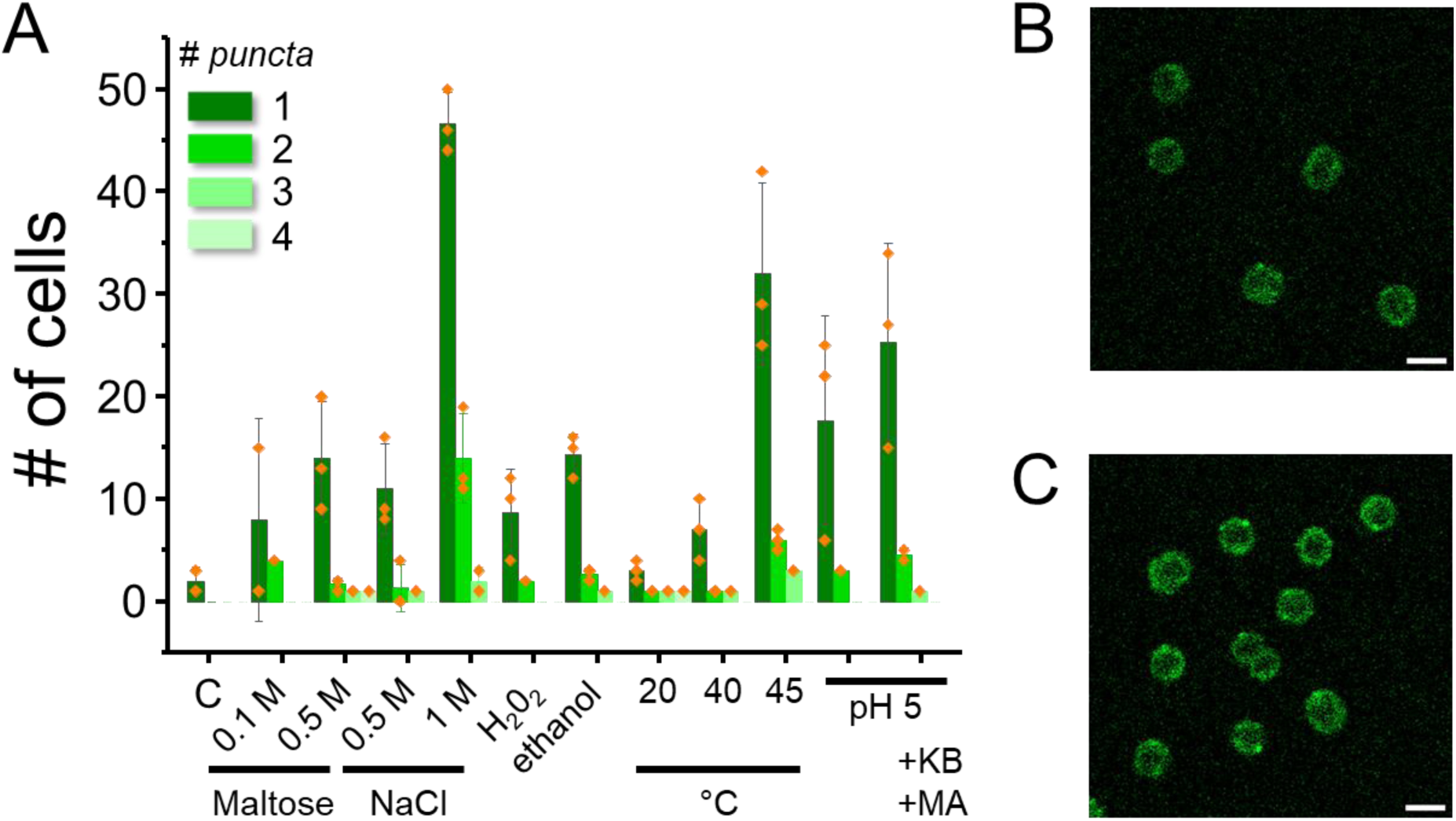
Stress-induced formation of IM30 *puncta* in *Synechocystis*. (A) *Synechocystis* cells expressing IM30–mVenus were subjected to the indicated stress conditions (see Methods for details). Fifteen minutes after stress application, cells were imaged and analyzed. The proportion of cells exhibiting *puncta*, as well as the average number of *puncta* per cell, was quantified. Representative images of stressed cells are shown in Figure S1. The individual results of three biological replicates with the statistical evaluation are shown in Figure S7. KB: potassium benzoate; MA: methylamine hydrochloride. (B) Fluorescence microscopy of *Synechocystis* cells expressing mVenus-tagged IM30 15 min after shifting the medium pH to 5.0. Acid stress triggers the rapid formation of IM30 *puncta*. (C) Cells after shifting the medium pH to 5.0 and the addition of 40 mM benzoate and 40 mM methylamine to collapse the ΔpH. Scale bar in (B + C): 2 µm.

For us, the observation that lowering the pH also drastically increased the formation of puncta was of special interest. Neutralophilic bacteria can typically grow well at environmental pH values from ∼5.5–9.0 and generally maintain their cytoplasmic pH in a narrow range of ∼7.5–7.7 (Padan et al., 2005; Slonczewski et al., 2009). However, lowering the pH of the medium will clearly affect metabolic processes (especially at the cytoplasmic membrane) and stress the cells. As we observed, *Synechocystis* cells form IM30 *puncta* in the cytoplasm in response to a more acidic external pH. This is of special physiological relevance, as cyanobacteria possess an additional internal membrane system, the TMs, in which a pH gradient is established (Mullineaux, 2014), and transient TM defects can allow proton leakage from the thylakoid lumen into the cytoplasm, resulting in a spatiotemporally lowered pH in the cytoplasm.

### pH-induced in vivo and in vitro formation of IM30 condensates

Acidic pH-induced condensate formation might be of special importance for TM systems, where a proton gradient is established across the TM (Belkin et al., 1987). At damaged TM sites, the local pH can easily drop to <6, and thus, the observation that lowering the pH triggers condensate formation is physiologically highly relevant. In the above-described experiment, we have lowered the external pH of the medium to 5 and monitored whether this triggered the formation of intracellular IM30 *puncta* (Figure 7A, B). However, while we indeed observed *puncta* formation in ∼20% of all cells, with most cells containing a single *punctum*, the cells clearly maintained their cytoplasmic pH. To overcome this buffering mechanism, at least to some extent, we next lowered the external pH again to 5.0 and added potassium benzoate and methylamine hydrochloride to collapse the transmembrane pH gradient (Martinez et al., 2012). In these cells, we observed increased formation of IM30 *puncta* (Figure 7A, C), indicating that lowering the pH triggers IM30 phase-separation and condensate formation *in vivo*, in living cyanobacterial cells.

Also, *in vitro*, the monomeric IM30* formed condensates already at a pH 6.0 in the absence of PEG and NaCl (Figure 8A), whereas at pH 7.5, NaCl and PEG were needed (Quarta et al., 2026). Turbidity measurements followed by differential interference contrast (DIC) microscopy enabled us to monitor condensate formation at varying pH and protein concentrations in the complete absence of salt, or any crowding agent (Figure 8C, D).

**Figure 8:**
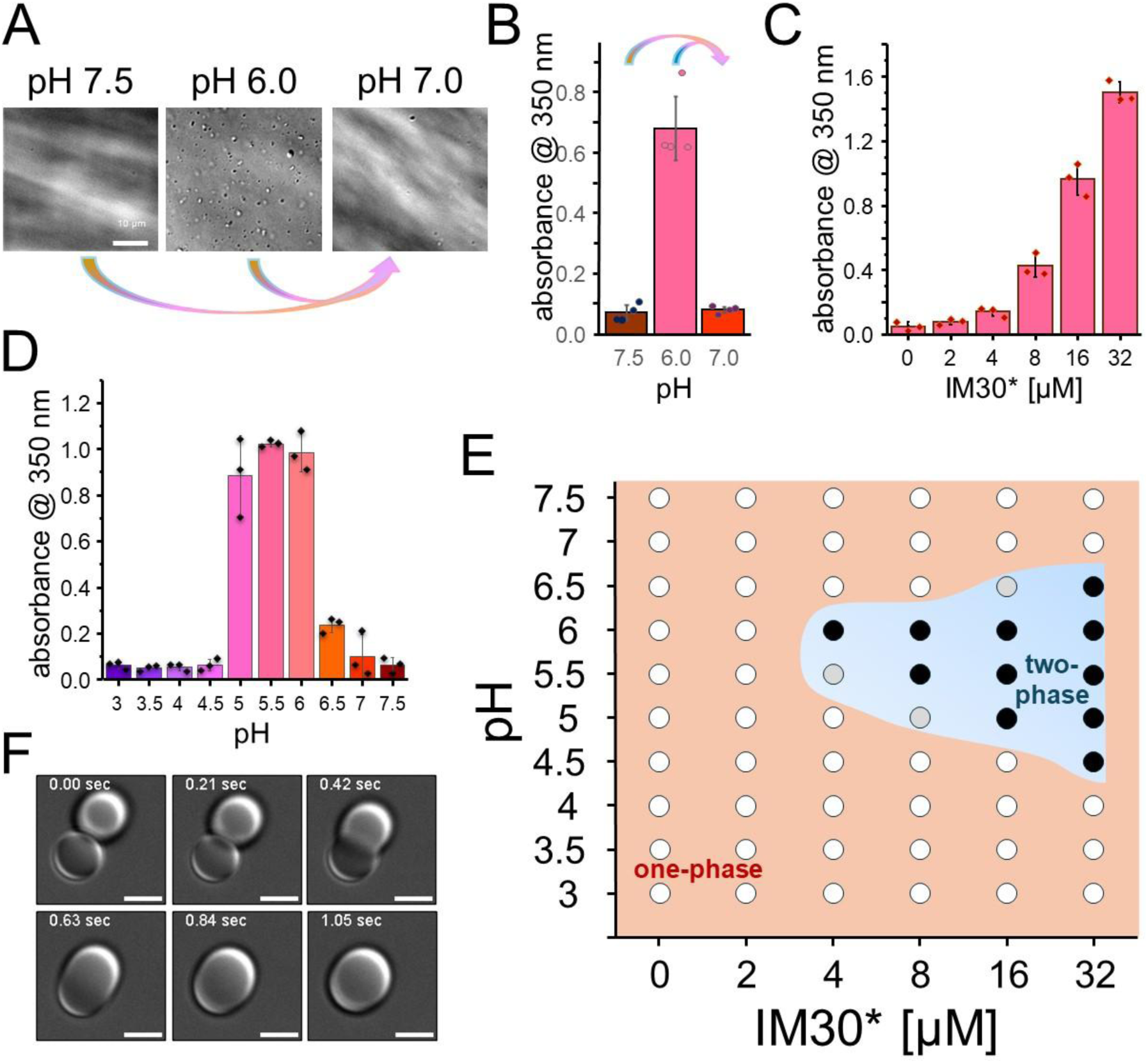
pH-induced formation of IM30* condensates. (A) DIC images of 32 µM IM30* at pH 7.5, pH 6.0 and a mixed solution with a final pH of ∼7.0. Representative images of four independent experiments are shown. Scale bar: 10 µm. (B) Turbidity of the IM30* solutions shown in (A) at different pH values. Error bars represent SD (n = 4 biological replicates). (C) Turbidity of IM30* solutions at a constant pH 5.5 and different protein concentrations in the absence of PEG and NaCl. (D) Turbidity of an 8 µM IM30* solution at different pH values. Error bars in (C-D) represent SD (n = 3 biological replicates). (E) Phase diagram of IM30* at various pH values and protein concentrations using turbidity measurements. At a turbidity above 0.16, condensates became observable via DIC; thus, this value was set as the lower limit for condensate formation (grey). Values ≥ 0.32 were defined as significant condensate formation (black). (F) DIC microscopy snapshots of a fusion event. The time elapsed after the condensates came into contact is indicated in each image. The scale bar is 3 µm. Condensates were formed at pH 5.5, in 10 mM HEPES and 10 mM P_i_-buffer. A video showing condensate fusion is available in the supplements (Movie S1).

At a constant protein concentration of 8 µM, we observed the formation of condensates via DIC microscopy, accompanied by an increase in turbidity, upon slightly lowering the pH to about 6.0 (Figure 8D). Further acidification below pH 5 resulted in the return to a single-phase regime and base levels of turbidity. Turbidity at a constant pH of 5.5 increased with increasing protein concentrations and indicated a c_sat_ in the low µM range (Figure 8C). Increasing the protein concentration above this critical concentration likely led to higher volume fractions of the condensed phase, while keeping the protein concentration in the dilute phases constant, as described for simple one-component systems (Alberti et al., 2019). Summarizing multiple turbidity measurements into a phase diagram (Figure 8E) revealed a re-entrant phase transition characterized by two phase transitions in response to pH, with a clear preference for the formation of condensates at a pH around 5.5-6.0, a value close to the calculated isoelectric point (pI) of IM30* (5.75). In fact, condensate formation in response to pH is predicted to typically occur around a protein’s pI (Adame-Arana et al., 2020).

Importantly, condensate formation induced by lowering the pH was reversible: when an IM30* solution at pH 6.0 (containing condensates) was mixed with an IM30* solution at pH 7.5 (where no LLPS was observed), the resulting mixture had a pH of 7.0 and condensates dissolved completely (Figure 8A, B). This clearly demonstrates that condensate formation is highly reversible, indicating a liquid nature of the condensates. In fact, we observed that two adjacent condensates can fuse into one single spherical condensate within a second (Figure 8F, Movie S1), plus condensates often wet the glass surface of the chamber during observation in DIC microscopy experiments after extended incubation periods (Figure S8). All observations show the fluid nature of the condensates.

To study the dynamic features of the pH-induced IM30* condensates in more detail, we followed the fluorescence recovery after photobleaching (FRAP) of single condensates (Figure 9A, B). Upon full-droplet bleaching, we observed rapid fluorescence recovery to ∼75% of the initial fluorescence after 1 min and a >90% fluorescence recovery after 10 min (Figure 9B). From the recovery curves, we extracted two key kinetic parameters: the time required for the fluorescence intensity to reach half of its final value (the recovery half-time, t₁/₂) and the apparent diffusion coefficient (D). Based on the analysis of six individual fluorescence recovery curves, the mean half-time was determined to be t₁_/_₂ = 14.8 ± 5.4 s, in close agreement with the value derived from the averaged curve (Figure 9B, 14.9 s). Using the individual half-times and corresponding illuminated areas, the mean diffusion coefficient was calculated as D = 0.093 ± 0.024 µm² s⁻¹, a value consistent with those typically reported for biomolecular condensates formed via LLPS.

**Figure 9:**
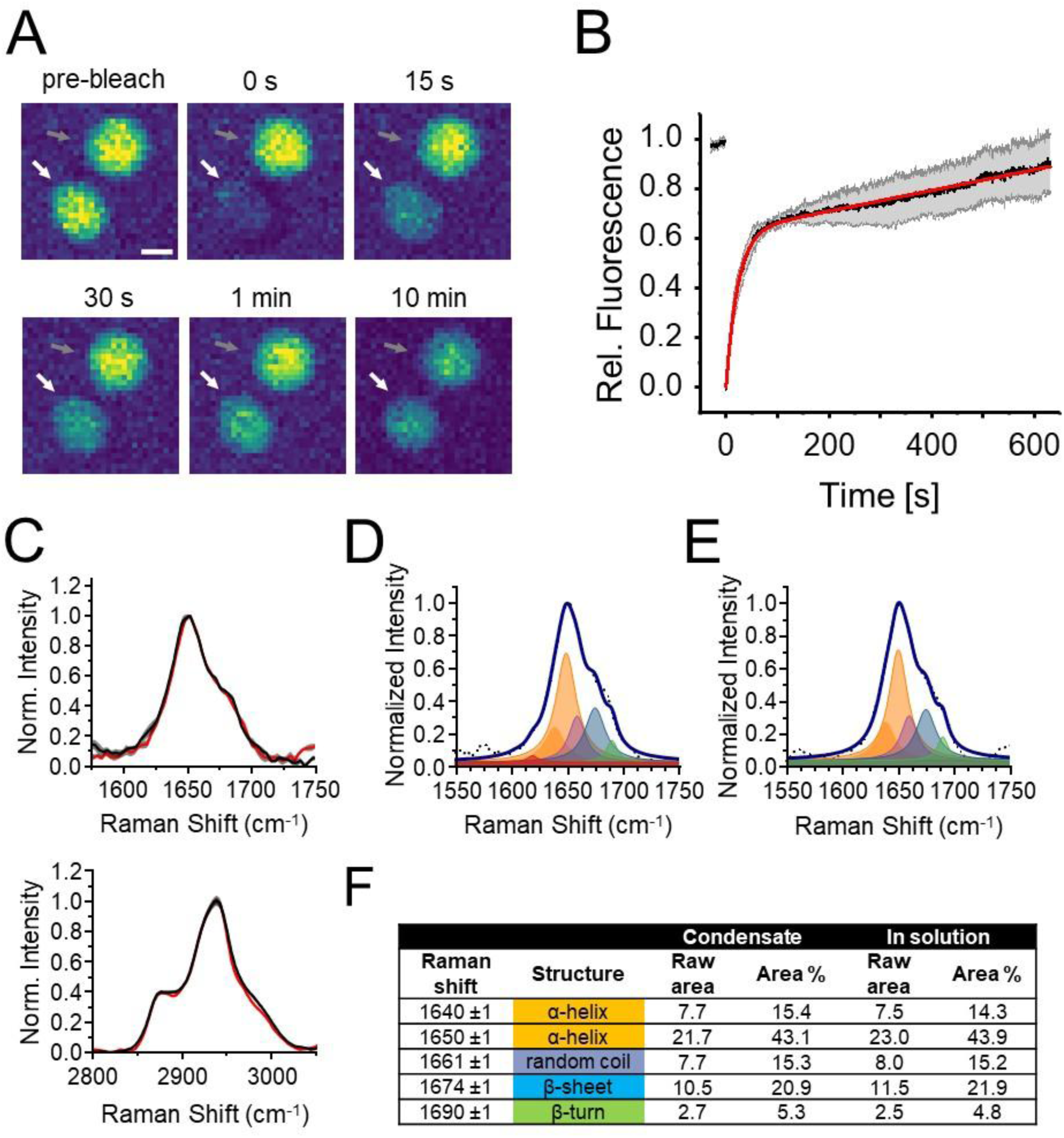
Structure and dynamics of IM30* condensates. (A) FRAP experiments: fluorescence microscopy images of a fully bleached (white arrow) and a non-bleached (gray arrow) pH-induced IM30* condensate before and after the bleach pulse. Non-bleached condensates were used as controls to correct for acquisition bleaching. The scale bar is 2 µm. (B) Normalized fluorescence intensity measured in FRAP experiments with pH-induced IM30* condensates. Error bars represent SD (n=3 biological replicates (N = 6 condensates)). The fit (red line) yielded a t_0.5_= 14.9 ± 0.1 s. (C) *Top*, BCARS in the Amide I region of IM30* in pH-induced condensate (black, pH 5.5) and liquid (pH 7.6, red) form. *Bottom*, BCARS in the CH regions of IM30* in pH-induced condensate (black, pH 5.5) and liquid (pH 7.6, red) form. (D, E) IM30* Amide I band deconvolution in condensate (D) and liquid (E) form. Proteins were analyzed in 10 mM HEPES buffer at pH 7.6 (free proteins in solution) and pH 5.5 (condensates). (F) Deconvolution of the IM30* Amide I region (compare D, E) using Lorentzian peaks assigned to specific secondary structures of proteins. A nearly identical secondary structural composition is obtained based on the areas. Condensates were formed at pH 5.5, in 10 mM HEPES and 10 mM P_i_-buffer.

To further classify the state of pH-induced IM30* condensates (fluid *vs*. gel-like), we next monitored the structure of IM30* monomers within the condensates and compared them to their free structure in solution. To this end, we employed broadband coherent anti-Stokes Raman scattering (BCARS) microspectroscopy to quantitatively assess potential structural changes (Figure 9C-F). A comparison of the Amide I (*ca*. 1600 to 1700 cm^-1^) and CH (2800 to 3100 cm^-1^) regions, which are associated with the C=O in the peptide bond and C-H stretching vibrations, respectively, revealed no discernible differences between the protein structure in condensates *vs*. in solution (Figure 9C). Furthermore, deconvolution of the Amide I region using Lorentzian peaks assigned to specific secondary structures of proteins yielded nearly identical secondary structure distributions based on the areas obtained (Figure 9D-F). Thus, the structure of the protein monomers remains unchanged when incorporated into a condensed assembly.

Together, these observations collectively confirm that pH is a direct, physiologically relevant trigger of IM30 condensate formation. Lowering the pH induces the reversible formation of liquid-like condensates. The structure of protein monomers does not change upon condensation *in vitro*, confirming phase separation occurs without conformational rearrangement, similar to what is seen for the mammalian FUS protein (Murthy et al., 2019), and likely via the stably structured α1-3 helical-hairpin (compare above). Combined with the dynamic, rapid exchange kinetics observed *in vivo*, these findings strongly suggest that IM30 condensate formation in living cells is finely tuned by the cellular pH landscape (besides other environmental changes).

### In vitro formation of IM30 wt condensates induced by oligomer disassembly

The structure of the IM30 barrels appears to inhibit condensate formation *in vitro* and leads to aggregate formation under conditions where IM30* readily forms condensates (Quarta et al., 2026). Previously, we have shown that destabilizing the IM30 wt oligomeric structure with urea allows condensate formation when both NaCl and PEG are added (Quarta et al., 2026). We therefore next pre-destabilized the wt IM30 barrel structure by adding 3 M urea, a condition under which the oligomer disassembles while the α1-3 helical hairpin motif remains stably folded (Quarta et al., 2024). We compared pH-induced condensate formation of this urea-destablilized IM30 and IM30* (Figure 10A, B). Turbidity measurements indicated a similar behavior for both pH-induced IM30* and urea-destablized IM30; however when combined with imaging, we see clear differences between the two protein systems.

**Figure 10:**
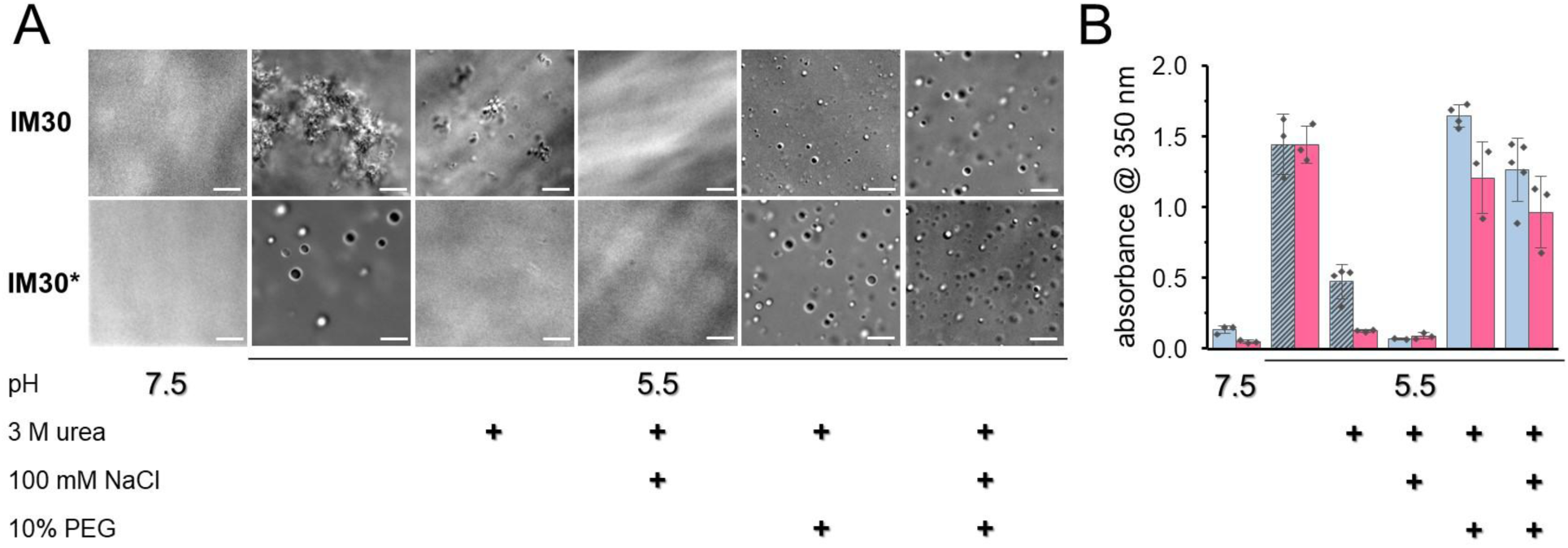
Phase behavior of IM30 and IM30 under varying physicochemical conditions. (A) DIC microscopy images of 32 μM IM30 or IM30* incubated at pH 7.5 and pH 5.5, in the absence or presence of 3 M urea, 100 mM NaCl, and/or 10% PEG, as indicated. (B) Bar graph depicting solution turbidity for 32 μM IM30 wt (blue bars) and IM30* (red bars) under the indicated buffer conditions. Diagonally striped bars indicate conditions under which large aggregates or particulate species formed, as opposed to well-defined liquid condensates. Notably, aggregation was observed exclusively with wt IM30. Data represent mean values ± SD from three independent biological replicates.

While IM30 failed to assemble into condensates at pH 5.5 in the presence of 3 M urea, it did form aggregates to a limited extent (Figure 10A, B). While the addition of 100 mM NaCl alone did not induce condensate formation, IM30 underwent phase separation and formed biomolecular condensates at pH 5.5, 3 M urea, when PEG was additionally present, as observed via DIC microscopy (Figure 10A). It appears that pH-induced IM30 aggregates arise because of attractive intermolecular electrostatic interactions that are maximized near the protein pI, where the protein net charge is zero. The formation of pH-induced aggregates may be suppressed in the presence of NaCl, likely due to charge screening that reduces electrostatic interactions and the simultaneous destabilizing effect of urea on hydrogen bonds.

In summary, by systematically screening the parameters that were previously identified to promote the formation of IM30* condensates (Quarta et al., 2026), we now show that differences in pH affect the formation of biomolecular IM30 assemblies. The formation of IM30 condensates appears to depend on a combination of at least pH and the oligomeric state (plus potentially other factors). Taken together, these data clearly show that oligomer disassembly enables the formation of IM30 wt condensates and that IM30* is a valid model for studying IM30 condensate formation in greater detail.

## Discussion

### Stress-induced IM30 puncta in vivo recapitulate in vitro condensates

Biomolecular condensates emerge through LLPS of protein solutions, yielding a dense phase in equilibrium with a dilute phase. Condensate assembly necessitates protein concentrations surpassing a critical saturation threshold concentration (c_sat_) and can be triggered by environmental perturbations, including alterations in pH, ionic strength, or macromolecular crowding, which modulate biopolymer-solvent or intermolecular interactions (Alberti and Dormann, 2019; Banani et al., 2017; Hyman et al., 2014; Shin and Brangwynne, 2017). These condensates typically exhibit liquid-like properties, facilitating swift molecular exchange between dense and dilute phases. Over time, certain spherical droplets may evolve into gel-like structures (Andre and Spruijt, 2020; Franzmann et al., 2018; Van Lindt et al., 2021), and such liquid-to-solid transitions can serve physiological roles (Guo et al., 2022) or, conversely, precipitate dysfunctional aggregates and toxicity (Alberti and Dormann, 2019; Dormann and Lemke, 2024; Mathieu et al., 2020; Ray et al., 2020).

While well-characterized in eukaryotes, condensate formation in bacteria remains largely underexplored (Sasazawa et al., 2024; Stevens and Lasker, 2025). In the cyanobacterium *Synechocystis* sp. PCC 6803, IM30 *puncta* arise under high-light or salt stress (Bryan et al., 2014; Quarta et al., 2026). Our findings now extend this to diverse stressors, positioning *puncta* as a generic stress-adaptive response. Although typically solitary under moderate stress, multiple *puncta* arise under severe conditions, suggesting a graded cellular reaction.

Although their precise *in vivo* composition was ambiguous, *puncta* have recently been associated with biomolecular condensates (Quarta et al., 2026). Live-cell imaging now demonstrates uniform cytoplasmic distribution of IM30 in *Synechocystis* in *E. coli* cells at subsaturating levels, while *puncta* manifested upon exceeding c_sat_, a defining LLPS trait. Notably, stress-induced *puncta* are present even below c_sat_, reinforcing an LLPS mechanism. Super-resolution microscopy confirmed the spherical morphology of *in vivo* IM30 structures, similar to that of condensates. FRAP analysis of IM30-mVenus fusions revealed considerable fluorescence recovery within *puncta*, indicative of a fluid core and dynamic IM30 exchange with the cytosol. Collectively, these properties, stress responsiveness, liquidity, and reversibility, establish IM30 *puncta* as *bona fide* biomolecular condensates.

While previous studies, in addition to the above observations, suggest that IM30 forms condensates *in vivo*, it has been speculated that the observed *puncta* represent clustered or aggregated IM30 oligomers (Bryan et al., 2014; Gates et al., 2022; Gutu et al., 2018). This hypothesis, however, is contradicted by two independent observations:

First, IM30*, an IM30 variant engineered to be deficient in oligomerization (Heidrich et al., 2016; Junglas et al., 2020), still generated *puncta* structures when expressed in living cyanobacterial cells (Figures 4, 6). The persistence of *puncta* in the absence of the canonical barrel-shaped oligomer thus demonstrates that the formation of such higher-order assemblies does not depend on the canonical IM30 oligomeric scaffold. Consequently, *puncta* formation cannot be explained solely as the accumulation of homo-oligomeric IM30 rings and/or rod assemblies. Second, the morphology of the *puncta* typically is spherical (Figure 4), which is inconsistent with an alternative model that envisions the structures as planar “membrane carpets” formed by IM30 coating the TM or cytoplasmic membrane (Junglas et al., 2025; Junglas et al., 2020). The isotropic shape, therefore, argues for a liquid-like condensate rather than a two-dimensional protein sheet.

A quantitative assessment of the IM30 intracellular concentration further substantiates the relevance of phase separation under physiological conditions. Recent measurements (Jackson et al., 2023) have established that a single *Synechocystis* cell contains, at minimum, 16 µM IM30. This value represents a conservative lower bound, because the calculation does not account for the substantial volume occupied by the TM system, which effectively reduces the cytoplasmic volume accessible to soluble proteins (Quarta et al., 2026). Crucially, our i*n vitro* phase-diagrams (Figure 8D) indicate that, when condensate formation is triggered by acidification, the c_sat_ of monomeric IM30 lies at approximately 4 µM. The intracellular concentration of IM30 therefore exceeds this critical threshold by a factor of 4 or more, placing the protein comfortably within the regime in which condensate formation is thermodynamically favored. This concordance between the *in vivo* concentration range and the *in vitro* determined c_sat_ aligns with the observed formation of IM30 *puncta* in living cells.

It should be emphasized, however, that c_sat_ is not a static parameter (Harmon et al., 2017; Hyman et al., 2014). Both *in vitro* and *in vivo*, it is modulated by a suite of physico-chemical variables, including temperature, ionic strength, pH, and potentially post-translational modifications. Accordingly, the precise value of IM30’s c_sat_ under native cellular conditions may differ from the experimentally derived 4 µM. Integrating these lines of evidence, the persistence of *puncta* in IM30* mutants, their spherical architecture, their fluid nature, and intracellular concentrations decisively surpassing c_sat_, we conclude that IM30 *puncta* constitute authentic biomolecular condensates rather than artifactual aggregates. This conclusion now establishes a coherent mechanistic framework for understanding IM30-mediated phase separation in cyanobacterial stress adaptation, highlighting its potential role as a conserved physiological response in prokaryotic systems.

### The helical hairpin formed by helices α1-3 triggers phase separation both in vitro and in vivo

*Puncta* formation has previously been documented for IM30 homologs in *Arabidopsis thaliana*, as well as for PspA in *Escherichia coli* and LiaH in *Bacillus subtilis*. In *A. thaliana*, IM30 overexpression yields diverse morphologies, including forked and web-like assemblies (Zhang et al., 2012), alongside *puncta* observed here and previously in *Synechocystis* (Bryan et al., 2014; Gates et al., 2022; Gutu et al., 2018). In contrast, *in vitro* condensates of *Synechocystis* IM30 wt and IM30* were predominantly spherical, without irregular shapes. Concordantly, live-cell imaging of IM30 or IM30* in *Synechocystis* primarily revealed spherical *puncta*, occasionally accompanied by small membrane-proximal patches (Figure 4) likely corresponding to previously described surface-covering carpets (Junglas et al., 2025; Junglas et al., 2020). Notably, IM30* and, to a greater extent, IM30 α0-3 also formed web-like condensates *in vivo* (Figure 6B). Although condensates are typically considered to be spherical, they can adopt non-spherical configurations, such as tubular networks (Emenecker et al., 2021; Fare et al., 2021; Jawerth et al., 2020; Linsenmeier et al., 2022; Mangiarotti et al., 2023). For instance, the TIS granules ER domain forms an interconnected tubular mesh around the endoplasmic reticulum (Ma and Mayr, 2018; Portz and Shorter, 2018). These morphologies may arise from sterically constrained interactions that orient truncated proteins to favor filaments or sheets over isotropic coalescence. In absence of the IM30 C-terminus, the α1-3 helical hairpins may engage in oriented interactions, promoting linear or sheet-like assemblies rather than compact droplets. This alignment could constrain condensates into mesh-like structures, particularly in cellular contexts. Additionally, polyanions such as RNA or DNA might bridge the cationic α1-3 hairpins, imprinting the condensate with the polymer’s contour instead of spherical compaction (Linsenmeier et al., 2022). Indeed, RNA binding to IM30 has recently been reported (Brenes-Alvarez et al., 2025), suggesting *in vivo* IM30 condensates may incorporate RNA or DNA.

In higher-order IM30 barrel oligomers, monomers forge multiple helix-helix contacts, with each monomer interfacing up to 16 others and α-helices forming an interwoven lattice via conserved interfaces (Gupta et al., 2021; Schlosser et al., 2023). Crucially, α1-3 hairpin interactions with neighboring hairpins underpin barrel assembly and stability. Here, we demonstrate that all C-terminally truncated IM30 variants retaining the α1-3 hairpin undergo *in vitro* phase separation, corroborated by *in silico* analyses indicating robust multivalent contacts within this domain sufficient for condensate formation. Structured helical hairpins can drive LLPS independently (Ramirez et al., 2024), as evidenced by proteins relying solely on such motifs (Basalla et al., 2023; Kozak and Kaksonen, 2022). Thus, α1-3 hairpin-mediated attractions initiate IM30 condensate formation. Consistent with this, IM30 α4-6 failed to phase separate *in vitro*, and IM30 α4-6-mVenus remained cytosolic in *Synechocystis*, akin to free mVenus diffusion. Although IM30 α1-3 (lacking α0) formed condensates *in vitro*, its mVenus fusion did not yield *puncta in vivo*, likely reflecting disparities in microenvironmental conditions and/or *c*_sat_. Helix α0 contacts α1-3 residues in IM30 oligomers (Gupta et al., 2021; Junglas et al., 2025; Liu et al., 2021), potentially enabling multivalency. Conversely, the disordered α4-6 region alone resisted phase separation under all tested conditions, whereas full-length IM30* exhibited enhanced propensity relative to IM30 α0-3, implying α4-6 modulation. In barrels, α4-6 form helices that interface multiple monomers, vital for oligomerization (Gupta et al., 2021; Junglas et al., 2025; Liu et al., 2021; Schlosser et al., 2023). Our *in silico* data suggest monomeric α4-6 residues access additional contacts, accounting for elevated phase separation (Figure 6C). This is consistent with expectations from polymer physics: both the strength of intermolecular interactions required for phase separation and the critical concentration c_sat_ needed to trigger it decrease as polymer chain lengths increase (Rubinstein and Colby, 2003). This suggests that α4-6, while nominally a poor solvent for itself, is simply too short to undergo phase separation on its own, a conclusion supported by experimental observations of apparent polymer exponents close to ½ for most IDPs (Tesei et al., 2024). Collectively, these findings indicate that α1-3 hairpin interactions predominantly drive *in vitro* and *in vivo* IM30 condensate formation as *puncta*. This motif, a conserved positively charged feature of prokaryotic as well as eukaryotic ESCRT-III proteins, implies a broadly applicable mechanism.

Future investigations should test whether α1-3-hairpin-driven LLPS is a conserved feature across different species. The mechanisms that underlie the observed formation of *in vivo puncta* by PspA in *Escherichia coli* and LiaH in *Bacillus subtilis* are likely analogous. In contrast to the prokaryotic proteins, eukaryotic ESCRT-III proteins can adopt a closed conformation when they are monomeric in solution, and the transition from the open, IM30-like state to the closed state involves extensive structural rearrangements of the α1-3 region (Tang et al. 2015; Bajorek et al. 2009; Lata et al. 2008; Henne et al. 2012). In its closed conformation, helix α5 of the yeast ESCRT-III protein Snf7 folds back onto the α1–α2 hairpin, thereby disrupting the structural continuity between helices α2 and α3 and partially occluding the helical hairpin. This rearrangement potentially hinders the formation of condensates.

### Physiological implications

IM30 resides in the cyanobacterial cytoplasm or chloroplast stroma, either as a soluble protein or bound to internal membranes (Bryan et al., 2014; Fuhrmann et al., 2009a; Gutu et al., 2018; Li et al., 1994). The cytoplasmic and thylakoid lumen pH in cyanobacteria grown at neutral pH are estimated at approximately 7.8 and 5.6, respectively (Belkin et al., 1987). Under physiological conditions, the cytoplasmic pH exceeds 7, placing soluble IM30 outside the pH range conducive to condensate formation identified here (Figure 8D). However, as established earlier, the *in vivo* IM30 concentration (at least 16 µM) substantially exceeds the critical saturation concentration c_sat_ (≈ 4 µM), priming IM30 for rapid phase separation upon localized perturbations. Proton accumulation at negatively charged membrane surfaces substantially lowers local pH (Yamashita and Voth, 2010), a phenomenon directly relevant to IM30, which binds anionic membranes (Hennig et al., 2015) and undergoes pH-dependent conformational changes that potentially enhance multivalency (Siebenaller et al., 2020). Photosynthetic light reactions generate a transmembrane pH (Belkin et al., 1987; Mullineaux, 2014), and TM damage, commonly induced by environmental stresses, allows proton efflux from the lumen into the cytosol. Consequently, sites of membrane damage experience sharp, localized pH drops (to ≤6.0) sufficient to activate IM30 phase separation, as confirmed by its enrichment at these regions (Gates et al., 2022). The pH threshold of 5.5–6.0 for phase separation determined herein is therefore not only physiologically attainable but optimally tuned for IM30 recruitment to compromised membranes. Biomolecular condensates often form under stress to safeguard cellular integrity, and IM30 condensates indeed emerge prominently *in vivo* during stresses that impair membrane stability, as here evidenced by *puncta* formation in *Synechocystis* under diverse conditions, suggesting that pH-mediated environmental cues directly trigger phase separation to regulate IM30 function. In this context, IM30 potentially acts as a rapid sensor and effector for membrane perturbations. Given IM30’s established role in membrane reshaping, its stress-induced condensates likely drive stabilization or repair processes. Critically, condensates enable the concentrated, transient accumulation of unassembled IM30 monomers, bypassing the energetic cost of disassembling preformed oligomers, a prerequisite for forming the previously observed membrane-bound assemblies, such as carpets or membrane-internalizing oligomers (Aseeva et al., 2004; Fuhrmann et al., 2009a; Gupta et al., 2021; Junglas et al., 2025; Junglas et al., 2020; Liu et al., 2021; Naskar et al., 2025; Pan et al., 2024; Saur et al., 2017; Schlösser et al., 2023; Siebenaller et al., 2019; Thurotte and Schneider, 2019).

### Limitations of the study/Open questions

This study strongly suggests that IM30 undergoes LLPS to form biomolecular condensates both *in vitro* and *in vivo*. However, LLPS and the formation of other higher-order IM30 assemblies, such as membrane-covering “carpets,” rings, and rods that drive membrane fission and fusion (Junglas et al., 2025; Junglas et al., 2020), require disassembly of preexisting large homo-oligomeric complexes. These assemblies potently modulate IM30’s membrane-binding activity, yet the even cytoplasmic distribution of IM30 observed here and elsewhere indicates that higher-order polymers are not predominant under basal conditions. In *Synechocystis* and *Chlamydomonas reinhardtii*, IM30 engages Hsp70 chaperones to dynamically regulate the monomer–oligomer equilibrium (Bryan et al., 2014; Liu et al., 2007), thereby ensuring ample free monomers for rapid condensate formation. The use of monomeric IM30* here effectively models this state, but wt IM30 likely maintains a finely tuned equilibrium among monomers, defined oligomers, and condensates, each fulfilling specialized roles, with proportions controlled by chaperones and other partners whose mechanisms merit elucidation.

IM30 condensates emerge robustly *in vivo* during environmental stresses, including rapid recruitment to photoinduced membrane damage sites (Gates et al., 2022). These observations, aligned with IM30’s supersaturation above and responsiveness to localized pH drops at damaged membranes, solidify LLPS as a physiological mechanism underpinning IM30’s membrane association. Nonetheless, the dynamic interplay between condensates, membrane remodeling, and overall cellular homeostasis demands further dissection to fully illuminate IM30’s protective functions.

## Methods

### Cloning, expression, and purification of IM30 variants for in vitro studies

Construction of the plasmids used to express the genes coding for *Synechocystis* sp. PCC 6803 *im30* and *im30\** has been described previously (Fuhrmann et al., 2009a; Junglas et al., 2020). The plasmids for expression of the truncated IM30 variants IM30_α0-3_ and IM30_α1-3_ were based on pRSET_IM30 and were constructed by introducing a stop codon after the base triplet coding for amino acid 157. Via Gibbson assembly, the sequence coding for *Syn*IM30 α1 (residues 1-26) or amino acids 1-156 were removed, resulting in pRSET_IM30_α1-3_, and pRSET_IM30_α4-6_, respectively. A Gly-Ser linker of seven amino acids was added after the IM30 C-terminus before the mVenus fluorescence tag, which has previously been shown to result in a functional fusion protein (Bryan et al., 2014). The sequences of all the plasmids were confirmed by sequencing (Eurofins, Ebersberg, Germany).

For *in vitro* analyses, all protein variants were expressed in *E. coli* BL21 (DE3) grown overnight in LB medium at 37 °C. Cells were harvested by centrifugation (3000 × g, 4 °C), resuspended in 20 mM imidazole purification buffer (300 mM NaCl, 20 mM imidazole, 50 mM phosphate, pH 7.6), and lysed by sonication at 4 °C. Cell debris was removed by centrifugation (12000 × g, 4 °C), and His-tagged proteins bound to the Ni-NTA columns were washed with increasing amounts of imidazole (20, 50, 100 mM). Proteins were eluted with a purification buffer containing 500 mM imidazole. The buffer was exchanged using PD-10 columns or dialysis, and the proteins were concentrated using centrifugal filters (Merck, Darmstadt, Germany) with molecular weight cut-offs of 30 kDa for IM30 and IM30-mVenus, 10 kDa for IM30* and IM30*-mVenus, and 3 kDa for the truncated IM30 variants IM30_α0-3_, IM30_α1-3_, and IM30_α4-6_. Protein concentrations were determined with a Bradford assay using bovine serum albumin (BSA) as a standard for the calibration curve, and the proteins were frozen in liquid nitrogen and stored at −20 °C until use. To validate the purity of aliquoted protein samples, proteins (1 µg/lane) were separated on a 12% SDS-PAGE-gel and stained after electrophoresis with Coomassie Brilliant Blue R250.

### Turbidity measurements and phase diagrams

Turbidity measurements were performed in 384-well plates (Cellvis, Mountain View, CA, USA) using an Omega plate reader (BMG LABTECH, Ortenberg, Germany). To maintain constant buffer concentrations at varying pH values, we used phosphate buffers at empirically determined ratios of Na_2_HPO_4_, NaH_2_PO_4_ and H_3_PO_4_ solutions, resulting in a final buffer concentration of 10 mM phosphate and 10 mM HEPES at the indicated pH values. 60 µL of protein samples with the specified concentrations were prepared in either phase separation buffer (10 mM phosphate, 10 mM HEPES, pH as indicated) or PEG/NaCl buffer (20 mM HEPES pH 7.6, 10% (w/v) PEG-8000, NaCl as indicated) at the indicated urea concentrations. To ensure well-mixed and rapidly equilibrated protein solutions, the proteins were pre-mixed with 4 M urea if applicable. The pre-mixed protein was then added to the phase separation buffer or PEG/NaCl buffer to reach the final composition for the experiment. 50 µL were transferred to a 384-well plate and incubated for five min. at room temperature. Turbidity, defined as the absorbance value at 350 nm, was determined in the range of 300–600 nm, 20 flashes per well, and a 2-nm step width at room temperature. The samples were subsequently imaged using DIC microscopy to distinguish condensates and aggregates. Data from three independent turbidity measurements were combined to visualize condensate-forming conditions in a phase diagram by setting a minimal measured turbidity value for the indication of phase separation to 0.13 and a second value at twice the minimal value to give an indication of the changes at the boundaries.

### Differential interference contrast and fluorescence microscopy

Samples were prepared as described above for the turbidity measurements and imaged after 15 min incubation time using an Axio Observer.Z1 (Carl Zeiss, Jena, Germany) equipped with a 63x oil objective in differential interference contrast (DIC) mode, 500 ms acquisition time and 4.5-5.5 V lamp voltage. Fluorescence images were acquired using 3 ms acquisition time, 20% power of a 475 nm LED light source for the mVenus channel. The images were evaluated using FIJI software (Schindelin et al., 2012) by automatically adjusting the brightness of the DIC and fluorescence images.

For fluorescence recovery after photobleaching (FRAP) experiments, sample chambers were assembled by lining two stripes of double-sided tape on a glass slide and covering it with a cover slide to form the chamber. Samples were prepared as described above for the turbidity measurements, with 90% unlabeled and 10% of the respective mVenus-labeled proteins. FRAP experiments were performed on a Leica SP5 confocal microscope using a FITC laser and 63x objective at 1 % laser power for imaging. The sample was allowed to equilibrate inside the chamber for 15 min after mixing, and the regions of interest were selected for full-bleaching and partial bleaching of the condensates. Image size was 41.33 µm x 41.33 µm with 128 x 128 pixels to enable fast frame rates of 0.06 s. Regions of interest (ROIs) were defined in the software to mark individual condensate areas for bleaching. The time series was set to 30 s of pre-bleach acquisition, followed by 100% laser power bleach pulses for 10 frames (0.6 s) and 10 min of post-bleach acquisition. The FRAP experiments were evaluated using FIJI software (Schindelin et al., 2012) by selecting and measuring the mean intensity over time in ROIs for the background, control condensate, and condensates that were bleached by the bleach pulse. For each experiment, the background signal was subtracted, and the control condensate was used to correct for unwanted photobleaching. Normalized intensities were calculated using the following equation:

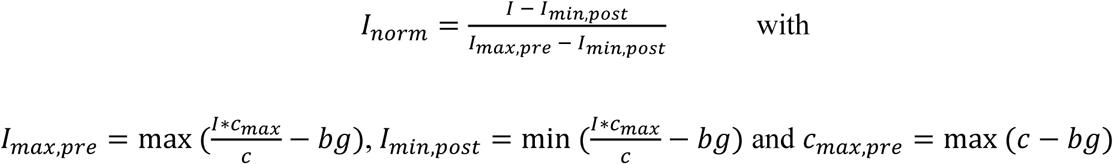

where *pre* and *post* refer to pre- and post-bleach time points being used for evaluation of extrema and *bg* and *c* are the mean intensities of ROIs from the background and the control condensate, respectively.

FRAP data were analyzed using ImageJ (Rueden et al., 2017). The normalized fluorescence intensities were fitted with a 2D diffusion model with a fixed boundary condition (Taylor et al., 2019). The resulting recovery curve was described by the equation:

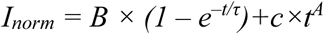

where *B* is the mobile fraction, and *τ* is the characteristic recovery time constant. The second terms takes into account the upward slope observed at longer times. The half-time of recovery (*t_1/2_*), *i.e.* the time it takes for fluorescence to recover to half its maximum, was then calculated as:

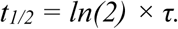

Assuming a uniform radius of the bleached area (*r*), the diffusion coefficient *D* was calculated based on the equation derived by Soumpasis (Soumpasis, 1983), which relates *t*_1/2_ to *D* by:

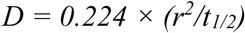

Since the bleached area (r²) was different for different experiments, for each individual experiment the diffusion coefficient was calculated from the fitted recovery time constant. Finally, the average and SD were calculated.

### Broadband Coherent Anti-Stokes Raman Scattering (BCARS) spectroscopy

Raman measurements were conducted using a custom-built broadband coherent anti-Stokes Raman scattering (BCARS) microscope, with comprehensive details available in the literature (Billecke et al., 2015). In summary, pump/probe and Stokes pulses were produced by a dual-output subnanosecond laser source (CARS-SM-30, Leukos). These pulses were then synchronously aligned in both space and time at the sample plane of an inverted microscope (Eclipse Ti-U, Nikon) and precisely focused onto the sample using a 0.85 NA air objective (LCPlan N, Olympus). After being separated from the excitation pulses, the BCARS signal was directed onto the slit of a spectrograph (Shamrock 303i, Andor Technology, Oxon, UK). The resulting scattered spectral components were captured using a cooled CCD camera (Newport DU920P-BR-DD, Andor Technology, Oxon, UK). Samples were mounted with a coverslip facing the collector and scanned using a piezo stage (Nano-PDQ 375 HS, Mad City Labs, Madison, USA) controlled by the LabView 2015 software (National Instruments, Austin, TX, USA). Moreover, the glass sample holders were chemically coated to enhance their hydrophobicity and condensate stability, if necessary, and sealed to prevent evaporation. The collected hyperspectral data were processed using IgorPro software (WaveMetrics, Lake Oswego, OR, USA). Raman-like spectra were obtained using a modified Kramer–Kronig transform (Liu et al., 2009). All spectra shown in this study were phase-retrieved using buffer alone, without macromolecules as a reference spectrum, and any background phase was eliminated using a Savitzky-Golay filter with a 2^nd^-order polynomial and a window size of approximately 400 cm^-1^. This method allows the acquisition and analysis of high-quality spectral data from the samples. The final spectra were obtained by averaging 10 spectra of each sample, with each spectrum based on a 2D mapping of 20 × 20 µm full of condensates or proteins in solution (1800 pixels or spectra). For deconvolution of the IM30* Amide I band of the monomers in the condensate form and free in solution, we employed a custom Python script after aligning the spectra using the Phe 1003 cm^-1^ *v*(C-C) Raman shift. This script was used to generate the initial parameter seeds for the Lorentzian peaks based on methodologies detailed previously (Chatterjee et al., 2022). Once these initial parameters were determined, they were fed into the peak analyzer function of OriginLab Pro (OriginLab Corporation, Northampton, MA, USA). This software performed further refinement, culminating in the final resolution of the peaks.

### IM30(*) condensate formation monitored in E. coli

The proteins IM30-mVenus, IM30*-mVenus, and free mVenus were expressed from pRSET-derived plasmids under the control of a T7 promoter, as described above. Plasmids were transformed into *E. coli* BL21(DE3) pLysE cells, and 20 mL overnight cultures were grown at room temperature with shaking at 130 rpm in LB medium supplemented with 100 µg/mL ampicillin and 30 µg/mL chloramphenicol. The following morning, a 20 mL culture was inoculated to an OD₆₀₀ of 0.1 using the overnight culture. Fluorescence emission of mVenus was monitored immediately after inoculation and hourly thereafter for four hours. Measurements were performed using a JASCO FP-8500 fluorescence spectrometer (JASCO Corporation, Tokyo, Japan). Emission spectra (500–570 nm) were recorded following excitation at 470 nm, with a scan rate of 100 nm/min, 1 nm step size, at a temperature of 25°C. Integration time was set to 0.5 s, and both excitation and emission slits were adjusted to achieve a spectral resolution of 2.5 nm.

Expression of the fusion protein and accumulation in the *E. coli* cytoplasm were visualized with a ZEISS Axio Observer.Z1 microscope (Zeiss, Oberkochen, Germany) equipped with a ZEISS ApoTome.2 to remove out-of-focus light, featuring a 63×/1.4 oil immersion objective. For image acquisition, cells were immobilized on 2% agarose. The mVenus was excited at 450-490 nm and imaged at 500-550 nm. Images were processed using the ZEN software (version 2.3.64.0) supplied by the vendor and ImageJ to adjust the contrast, merge the phase contrast and fluorescence channel and include the scale bars.

### Localization of fluorescently labeled IM30 variants in Synechocystis

The construction of *Synechocystis* sp. PCC 6803 cells stably expressing mVenus-tagged IM30 and IM30* proteins upon induction were described recently (Schlösser et al., 2025). To express truncated mVenus-labeled IM30 variants in living *Synechocystis* cells, the IM30 variants (as described above) containing the 7GS linker before the mVenus tag at the C-terminus of the protein were cloned into the plasmid pCK306, which allowed integration of the chimeric genes into a neutral site of the *Synechocystis* genome (*ssl0410* locus) and enabled rhamnose-inducible expression of chimeric genes controlled by the *E. coli rhaBAD* promoter (Kelly et al., 2018). *Synechocystis* wt cells were transformed with the pCK306-based plasmids, and kanamycin-resistant mutant cells were selected and passaged on BG11 plates containing increasing kanamycin concentrations. Complete segregation was confirmed by PCR.

For *in vivo* localization and 3D rendering of fluorescently labeled IM30, *Synechocystis* cells were grown in a shaker under constant warm white light (30 μmol⋅photons m^−2^⋅s^−1^) in BG11-medium (Rippka et al., 1979) supplemented with 5 mM glucose and 100 µg/ml kanamycin. Before the experiments, cells were diluted to an OD_750_ = 0.1 and 1 mg/ml L-rhamnose was added to induce gene expression. After two days (OD_750_ = 0.4-0.6), cells were imaged using a ZEISS Elyra 7 with Lattice SIM^2^ super-resolution microscope (Zeiss, Oberkochen, Germany) equipped with a 63×/1.4 oil objective, using a two-track configuration, a 642 nm laser for chlorophyll, and a 488 nm laser for mVenus excitation, with the filter set to SBS LP 560 and simultaneous imaging by two cameras. 13 phase images were captured as 1280 x 1280 pixels at 16 bits and super-resolution images were achieved by SIM^2^ reconstruction using the Zen software. For 3D rendering, raw data was collected at 0.110 μm slice intervals and processed using the SIM^2^ leap model, and 3D rendering was performed in Imaris (version 10.1.1). Images for this study were further processed using ImageJ (Rueden et al., 2017) to set the saturation levels and include scale bars.

### Monitoring stress-induced IM30 condensate formation in Synechocystis

*Synechocystis* sp. PCC cells expressing mVenus-tagged IM30 were cultured in a shaker at 30°C under continuous, low-intensity warm white light (30 µmol photons m^-2^ s^-1^) in BG11 medium (Rippka et al., 1979). To investigate the behavior of IM30 proteins *in vivo*, in living cyanobacterial cells, gene expression was gradually induced by adding increasing concentrations of L-rhamnose (0.05–1.0 mg/mL). 24 h after induction, cells were immobilized on a 2% agarose pad and visualized with a Zeiss Axio Observer.Z1 microscope (Zeiss, Oberkochen, Germany) equipped with a Zeiss ApoTome.2 using a 63x/1.4 oil immersion objective at room temperature. mVenus was excited at 450-490 nm and imaged at 500-550 nm. Images were processed using the ZEN software (version 2.3.64.0) supplied by the vendor and ImageJ to subtract the background, adjust the contrast and include the scalebars.

To study the effects of osmotic, pH, and temperature stress on IM30 behavior, gene expression was induced by the addition of 0.1 mg/mL L-rhamnose to cultures having an OD_750_ of ∼1.0. 24 h after induction, cultures were exposed to the respective stressors (see main text) prior to immobilization and imaging. For osmotic and pH stress, 200 µL of culture was centrifuged at 2000 g for 2 min; the supernatant was discarded, and cells were resuspended in BG11 medium containing the stressor. For pH stress, BG11 was buffered with acetic acid–acetate to pH 5. 0.25 mM H_2_O_2_ or 5 % ethanol (final concentrations), respectively, were directly added to the growth medium. Cells were incubated under stress conditions for 15 min at 30°C prior to imaging. For temperature stress, cells were incubated at the indicated temperature for 30 min prior to analysis.

### Fluorescence Recovery after Photobleaching (FRAP) measurements in salt-stressed Synechocystis cells

Salt stress was induced by adding 1 M NaCl to the growth medium, as previously described (Quarta et al., 2026). FRAP measurements were performed on immobilized cells using a Zeiss LSM980 laser scanning microscope (Carl Zeiss, Oberkochen, Germany), equipped with a 63×/1.4 oil immersion objective, at room temperature. mVenus was excited with a 514 nm laser at 1% power, with a master gain of 750 V and a 52 µm pinhole. Fluorescence emission was monitored between 517 and 570 nm. The time-lapse acquisition was set to 5 minutes, with images captured at 5-second intervals. Photobleaching of the target *puncta* was performed immediately after the first image, using 10% laser power of the 514 nm line. The FRAP experiments were analyzed as described above for the *in vitro*-formed condensates, with the exception that the additional term accounting for the slope (c × tA) was omitted.

### Contact Map Analysis

Coarse-grained (CG) molecular dynamics simulations were performed using the CALVADOS-2 force field, in which each amino acid was represented as a single bead and the solvent was treated implicitly (Tesei and Lindorff-Larsen, 2022). Initial system configurations were generated using HOOBAS (Girard et al., 2019). Electrostatic screening was incorporated by explicitly setting the inverse Debye length (κ) to 0.097 Å⁻¹, corresponding to an experimental salt concentration of 88.3mM. Simulations for both IM30 and IM30* were performed at 220 K, at which a distinct two-phase system forms, as indicated by the phase diagram reported in our previous work (Quarta et al., 2026). Simulations were performed using HOOMD-Blue (Anderson et al., 2020; Howard et al., 2019) in a slab geometry with dimensions of 3000 × 250 × 250 Å³, containing 216 protein chains. The system was evolved for 5 μs in the canonical (NVT) ensemble using Langevin dynamics, with an integration time step of Δt = 0.01τ, where τ corresponds to 10 fs. Trajectory data were recorded as 5000 snapshots per simulation, with the first 500 snapshots discarded to ensure equilibration. Simulation data management and workflow automation were performed using signac and signac-flow (Adorf et al., 2018; Dice et al., 2021; Ramasubramani et al., 2018).

Average inter-chain contact maps were computed from the simulation trajectories. A residue–residue contact C_ij_ was assigned a value of 1 when the distance between beads corresponding to residues i and j satisfied, ∣r_i_−r_j_ ∣ <σ_ij_, and 0 otherwise. The contact cutoff was defined as σij=1.5(σ_i_+σ_j_)/2, where σ_i_ and σ_j_ are the residue-specific bead sizes defined in the CALVADOS-2 force field. The contact frequency between each residue pair was computed by averaging Cij over all snapshots in the equilibrated trajectory.

## Data availability

Any additional information required to reanalyze the data reported in this paper is available from the lead contact upon request.

## Acknowledgements

This work was funded by the Max-Planck Graduate Center at the Max Planck institutes and the University of Mainz, as well as by the Deutsche Forschungsgemeinschaft (DFG, SCHN 690/16-1 to D.S. and CRC1551 (project no. 464588647) to M.G, M.B and D.S). Work in the lab of L.-N. L was supported by the Biotechnology and Biological Sciences Research Council (BB/W001012/1, BB/V009729/1, and BB/Y01135X/1 to L.-N.L.), the Royal Society (URF\R\180030 to L.-N.L.), and the Liverpool-Chinese Scholarship Council PhD studentship (X.G.). S.H.P. acknowledges support from the DFG (PA2526/3-1/2), Welch Foundation (F-2008-20220331) and National Science Foundation (#2146549).

## Author Contributions

**Ndjali Quarta:** Conceptualization, Data Curation, Formal Analysis, Investigation (molecular biology, protein expression and purification, turbidity and FRAP measurements, DIC micrcoscopy and low-resolution fluorescence microscopy), Methodology, Validation, Visualization, Writing – Original Draft. **Katrin Debrich:** Conceptualization, Data Curation, Formal Analysis, Investigation (*in vivo* phase separation), Methodology, Validation, Visualization, Writing – Original Draft. **Nadja Hellmann:** Conceptualization, Data Curation, Formal Analysis, Validation, Visualization, Writing – Original Draft, Writing – Review & Editing. **Xingwu Ge:** Investigation (high resolution fluorescence microscopy measurements). **Pablo G. Argudo:** Investigation (BCARS experiments), Writing – Original Draft. **Tika Ram Bhandari:** Formal Analysis, Investigation (CG simulations), Methodology, Validation, Visualization, Writing – Original Draft. **Mischa Bonn:** Funding Acquisition, Project Administration, Supervision, Writing – Original Draft, Writing – Review & Editing. **Martin Girard:** Methodology, Formal Analysis, Funding Acquisition, Project Administration, Supervision, Writing – Original Draft, Writing – Review & Editing. **Sapun H. Parekh:** Formal Analysis (BCARS experiments), Writing – Original Draft. **Lu-Ning Liu:** Formal Analysis (high resolution fluorescence microscopy measurements), Funding Acquisition, Supervision, Writing – Original Draft, Writing – Review & Editing. **Dirk Schneider:** Conceptualization, Formal Analysis, Methodology, Funding Acquisition, Project Administration, Resources, Supervision, Writing – Original Draft, Writing – Review & Editing.

## Materials & Correspondence

Correspondence and material requests should be addressed to D.S..

## References

Adame-Arana, O., Weber, C.A., Zaburdaev, V., Prost, J., and Julicher, F. (2020). Liquid Phase Separation Controlled by pH. Biophys J 119, 1590–1605.

Adorf, C.S., Dodd, P.M., Ramasubramani, V., and Glotzer, S.C. (2018). Simple data and workflow management with the signac framework. Comp Mater Sci 146, 220–229.

Al-Husini, N., Tomares, D.T., Pfaffenberger, Z.J., Muthunayake, N.S., Samad, M.A., Zuo, T., Bitar, O., Aretakis, J.R., Bharmal, M.M., Gega, A., et al. (2020). BR-Bodies Provide Selectively Permeable Condensates that Stimulate mRNA Decay and Prevent Release of Decay Intermediates. Molecular Cell 78, 670–682 e678.

Alberti, S., and Dormann, D. (2019). Liquid-Liquid Phase Separation in Disease. Annu Rev Genet 53, 171–194.

Alberti, S., Gladfelter, A., and Mittag, T. (2019). Considerations and Challenges in Studying Liquid-Liquid Phase Separation and Biomolecular Condensates. Cell 176, 419–434.

Alberti, S., and Hyman, A.A. (2021). Biomolecular condensates at the nexus of cellular stress, protein aggregation disease and ageing. Nat Rev Mol Cell Biol 22, 196–213.

Anderson, J.A., Glaser, J., and Glotzer, S.C. (2020). HOOMD-blue: A Python package for high-performance molecular dynamics and hard particle Monte Carlo simulations. Comp Mater Sci 173.

Andre, A.A.M., and Spruijt, E. (2020). Liquid-Liquid Phase Separation in Crowded Environments. Int J Mol Sci 21.

Aseeva, E., Ossenbühl, F., Eichacker, L.A., Wanner, G., Soll, J., and Vothknecht, U.C. (2004). Complex formation of Vipp1 depends on its alpha-helical PspA-like domain. J Biol Chem 279, 35535–35541.

Azaldegui, C.A., Vecchiarelli, A.G., and Biteen, J.S. (2021). The emergence of phase separation as an organizing principle in bacteria. Biophys J 120, 1123–1138.

Babst, M., Wendland, B., Estepa, E.J., and Emr, S.D. (1998). The Vps4p AAA ATPase regulates membrane association of a Vps protein complex required for normal endosome function. EMBO J 17, 2982–2993.

Banani, S.F., Lee, H.O., Hyman, A.A., and Rosen, M.K. (2017). Biomolecular condensates: organizers of cellular biochemistry. Nat Rev Mol Cell Biol 18, 285–298.

Banjade, S., Shah, Y.H., Tang, S., and Emr, S.D. (2021). Design principles of the ESCRT-III Vps24-Vps2 module. eLife 10.

Basalla, J.L., Mak, C.A., Byrne, J.A., Ghalmi, M., Hoang, Y., and Vecchiarelli, A.G. (2023). Dissecting the phase separation and oligomerization activities of the carboxysome positioning protein McdB. eLife 12.

Belkin, S., Mehlhorn, R.J., and Packer, L. (1987). Proton gradients in intact cyanobacteria. Plant Physiol 84, 25–30.

Billecke, N., Bosma, M., Rock, W., Fleissner, F., Best, G., Schrauwen, P., Kersten, S., Bonn, M., Hesselink, M.K., and Parekh, S.H. (2015). Perilipin 5 mediated lipid droplet remodelling revealed by coherent Raman imaging. Integrative biology: quantitative biosciences from nano to macro 7, 467–476.

Borcherds, W., Bremer, A., Borgia, M.B., and Mittag, T. (2021). How do intrinsically disordered protein regions encode a driving force for liquid-liquid phase separation? Curr Opin Struct Biol 67, 41–50.

Brangwynne, C.P., Eckmann, C.R., Courson, D.S., Rybarska, A., Hoege, C., Gharakhani, J., Julicher, F., and Hyman, A.A. (2009). Germline P granules are liquid droplets that localize by controlled dissolution/condensation. Science 324, 1729–1732.

Brenes-Alvarez, M., Ropp, H.R., Papagiannidis, D., Potel, C.M., Stein, F., Scholz, I., Steglich, C., Savitski, M.M., Vioque, A., Muro-Pastor, A.M., et al. (2025). R-DeeP/TripepSVM identifies the RNA-binding OB-fold-like protein PatR as regulator of heterocyst patterning. Nucleic Acids Res 53.

Bryan, S.J., Burroughs, N.J., Shevela, D., Yu, J., Rupprecht, E., Liu, L.N., Mastroianni, G., Xue, Q., Llorente-Garcia, I., Leake, M.C., et al. (2014). Localisation and interactions of the Vipp1 protein in cyanobacteria. Mol Microbiol 94, 1179–1195.

Brzezinski, M., Argudo, P.G., Scheidt, T., Yu, M., Hosseini, E., Kaltbeitzel, A., Lemke, E.A., Michels, J.J., and Parekh, S.H. (2024). Protein-Specific Crowding Accelerates Aging in Protein Condensates. Biomacromolecules 26, 2060–2075.

Buchkovich, N.J., Henne, W.M., Tang, S., and Emr, S.D. (2013). Essential N-terminal insertion motif anchors the ESCRT-III filament during MVB vesicle formation. Dev Cell 27, 201–214.

Carlton, J.G., and Baum, B. (2023). Roles of ESCRT-III polymers in cell division across the tree of life. Curr Opin Cell Biol 85, 102274.

Cermakova, K., and Hodges, H.C. (2023). Interaction modules that impart specificity to disordered protein. Trends in Biochemical Sciences 48, 477–490.

Chang, L.W., Lytle, T.K., Radhakrishna, M., Madinya, J.J., Velez, J., Sing, C.E., and Perry, S.L. (2017). Sequence and entropy-based control of complex coacervates. Nat Commun 8, 1273.

Chatterjee, S., Kan, Y., Brzezinski, M., Koynov, K., Regy, R.M., Murthy, A.C., Burke, K.A., Michels, J.J., Mittal, J., Fawzi, N.L., et al. (2022). Reversible Kinetic Trapping of FUS Biomolecular Condensates. Advanced science 9, e2104247.

Christ, L., Raiborg, C., Wenzel, E.M., Campsteijn, C., and Stenmark, H. (2017). Cellular Functions and Molecular Mechanisms of the ESCRT Membrane-Scission Machinery. Trends Biochem Sci 42, 42–56.

Das, R.K., and Pappu, R.V. (2013). Conformations of intrinsically disordered proteins are influenced by linear sequence distributions of oppositely charged residues. Proc Natl Acad Sci U S A 110, 13392–13397.

Dice, B.D., Butler, B.L., Ramasubramani, V., Travitz, A., Henry, M.M., Ojha, H., Wang, K.L., Adorf, C.S., Jankowski, E., and Glotzer, S.C. (2021). signac: Data Management and Workflows for Computational Researchers. Paper presented at: 20th Python in Science Conference (Texas).

Dominguez-Escobar, J., Wolf, D., Fritz, G., Hofler, C., Wedlich-Soldner, R., and Mascher, T. (2014). Subcellular localization, interactions and dynamics of the phage-shock protein-like Lia response in Bacillus subtilis. Mol Microbiol 92, 716–732.

Dormann, D., and Lemke, E.A. (2024). Adding intrinsically disordered proteins to biological ageing clocks. Nat Cell Biol 26, 851–858.

Elbaum-Garfinkle, S., Kim, Y., Szczepaniak, K., Chen, C.C., Eckmann, C.R., Myong, S., and Brangwynne, C.P. (2015). The disordered P granule protein LAF-1 drives phase separation into droplets with tunable viscosity and dynamics. Proc Natl Acad Sci U S A 112, 7189–7194.

Emenecker, R.J., Holehouse, A.S., and Strader, L.C. (2021). Sequence determinants of in cell condensate morphology, dynamics, and oligomerization as measured by number and brightness analysis. Cell communication and signaling: CCS 19, 65.

Engl, C., Jovanovic, G., Lloyd, L.J., Murray, H., Spitaler, M., Ying, L., Errington, J., and Buck, M. (2009). In vivo localizations of membrane stress controllers PspA and PspG in Escherichia coli. Mol Microbiol 73, 382–396.

Fare, C.M., Villani, A., Drake, L.E., and Shorter, J. (2021). Higher-order organization of biomolecular condensates. Open biology 11, 210137.

Franzmann, T.M., Jahnel, M., Pozniakovsky, A., Mahamid, J., Holehouse, A.S., Nuske, E., Richter, D., Baumeister, W., Grill, S.W., Pappu, R.V., et al. (2018). Phase separation of a yeast prion protein promotes cellular fitness. Science 359.

Fuhrmann, E., Bultema, J.B., Kahmann, U., Rupprecht, E., Boekema, E.J., and Schneider, D. (2009a). The vesicle-inducing protein 1 from Synechocystis sp. PCC 6803 organizes into diverse higher-ordered ring structures. Mol Biol Cell 20, 4620–4628.

Fuhrmann, E., Gathmann, S., Rupprecht, E., Golecki, J., and Schneider, D. (2009b). Thylakoid membrane reduction affects the photosystem stoichiometry in the cyanobacterium Synechocystis sp. PCC 6803. Plant Physiol 149, 735–744.

Galvanetto, N., Ivanovic, M.T., Del Grosso, S.A., Chowdhury, A., Sottini, A., Nettels, D., Best, R.B., and Schuler, B. (2025). Material properties of biomolecular condensates emerge from nanoscale dynamics. Proc Natl Acad Sci U S A 122, e2424135122.

Gates, C., Hill, N.C., Dahlgren, K., and Cameron, J.C. (2022). Kinetics and targeting of Vipp1 aggregation in cyanobacteria. BioRxiv.

Girard, M., Ehlen, A., Shakya, A., Bereau, T., and de la Cruz, M.O. (2019). Hoobas: A highly object-oriented builder for molecular dynamics. Comp Mater Sci 167, 25–33.

Guo, Q., Zou, G., Qian, X., Chen, S., Gao, H., and Yu, J. (2022). Hydrogen-bonds mediate liquid-liquid phase separation of mussel derived adhesive peptides. Nat Commun 13, 5771.

Gupta, A., Joshi, A., Arora, K., Mukhopadhyay, S., and Guptasarma, P. (2023). The bacterial nucleoid-associated proteins, HU and Dps, condense DNA into context-dependent biphasic or multiphasic complex coacervates. J Biol Chem 299, 104637.

Gupta, T.K., Klumpe, S., Gries, K., Heinz, S., Wietrzynski, W., Ohnishi, N., Niemeyer, J., Spaniol, B., Schaffer, M., Rast, A., et al. (2021). Structural basis for VIPP1 oligomerization and maintenance of thylakoid membrane integrity. Cell 184, 3643–3659 e3623.

Gutu, A., Chang, F., and O’Shea, E.K. (2018). Dynamical localization of a thylakoid membrane binding protein is required for acquisition of photosynthetic competency. Mol Microbiol 108, 16–31.

Harmon, T.S., Holehouse, A.S., Rosen, M.K., and Pappu, R.V. (2017). Intrinsically disordered linkers determine the interplay between phase separation and gelation in multivalent proteins. eLife 6.

Heidrich, J., Wulf, V., Hennig, R., Saur, M., Markl, J., Sonnichsen, C., and Schneider, D. (2016). Organization into Higher Ordered Ring Structures Counteracts Membrane Binding of IM30, a Protein Associated with Inner Membranes in Chloroplasts and Cyanobacteria. J Biol Chem 291, 14954–14962.

Henne, W.M., Buchkovich, N.J., and Emr, S.D. (2011). The ESCRT pathway. Dev Cell 21, 77–91.

Hennig, R., Heidrich, J., Saur, M., Schmuser, L., Roeters, S.J., Hellmann, N., Woutersen, S., Bonn, M., Weidner, T., Markl, J., et al. (2015). IM30 triggers membrane fusion in cyanobacteria and chloroplasts. Nat Commun 6, 7018.

Henry, J.T., and Crosson, S. (2013). Chromosome replication and segregation govern the biogenesis and inheritance of inorganic polyphosphate granules. Mol Biol Cell 24, 3177–3186.

Hess, N., and Joseph, J.A. (2025). Structured protein domains enter the spotlight: modulators of biomolecular condensate form and function. Trends in Biochemical Sciences 50, 206–223.

Hoang, Y., Azaldegui, C.A., Dow, R.E., Ghalmi, M., Biteen, J.S., and Vecchiarelli, A.G. (2024). An experimental framework to assess biomolecular condensates in bacteria. Nat Commun 15, 3222.

Hofmann, S., Kedersha, N., Anderson, P., and Ivanov, P. (2021). Molecular mechanisms of stress granule assembly and disassembly. Biochimica et biophysica acta Molecular cell research 1868, 118876.

Howard, M.P., Statt, A., Madutsa, F., Truskett, T.M., and Panagiotopoulos, A.Z. (2019). Quantized bounding volume hierarchies for neighbor search in molecular simulations on graphics processing units. Comp Mater Sci 164, 139–146.

Huber, S.T., Mostafavi, S., Mortensen, S.A., and Sachse, C. (2020). Structure and assembly of ESCRT-III helical Vps24 filaments. Science advances 6, eaba4897.

Hudina, E., Schott-Verdugo, S., Junglas, B., Kutzner, M., Ritter, I., Hellmann, N., Schneider, D., Gohlke, H., and Sachse, C. (2025). The bacterial ESCRT-III PspA rods thin lipid tubules and increase membrane curvature through helix alpha0 interactions. Proc Natl Acad Sci U S A 122, e2506286122.

Hyman, A.A., Weber, C.A., and Julicher, F. (2014). Liquid-liquid phase separation in biology. Annu Rev Cell Dev Biol 30, 39–58.

Jackson, P.J., Hitchcock, A., Brindley, A.A., Dickman, M.J., and Hunter, C.N. (2023). Absolute quantification of cellular levels of photosynthesis-related proteins in Synechocystis sp. PCC 6803. Photosynth Res 155, 219-245.

Janissen, R., Arens, M.M.A., Vtyurina, N.N., Rivai, Z., Sunday, N.D., Eslami-Mossallam, B., Gritsenko, A.A., Laan, L., de Ridder, D., Artsimovitch, I., et al. (2018). Global DNA Compaction in Stationary-Phase Bacteria Does Not Affect Transcription. Cell 174, 1188–1199 e1114.

Jawerth, L., Fischer-Friedrich, E., Saha, S., Wang, J., Franzmann, T., Zhang, X.J., Sachweh, J., Ruer, M., Ijavi, M., Saha, S., et al. (2020). Protein condensates as aging Maxwell fluids. Science 370, 1317-+.

Jin, X., Lee, J.E., Schaefer, C., Luo, X., Wollman, A.J.M., Payne-Dwyer, A.L., Tian, T., Zhang, X., Chen, X., Li, Y., et al. (2021). Membraneless organelles formed by liquid-liquid phase separation increase bacterial fitness. Science advances 7, eabh2929.

Junglas, B., Axt, A., Siebenaller, C., Sonel, H., Hellmann, N., Weber, S.A.L., and Schneider, D. (2022). Membrane destabilization and pore formation induced by the Synechocystis IM30 protein. Biophys J 121, 3411–3421.

Junglas, B., Huber, S.T., Heidler, T., Schlosser, L., Mann, D., Hennig, R., Clarke, M., Hellmann, N., Schneider, D., and Sachse, C. (2021). PspA adopts an ESCRT-III-like fold and remodels bacterial membranes. Cell 184, 3674–3688 e3618.

Junglas, B., Kartte, D., Kutzner, M., Hellmann, N., Ritter, I., Schneider, D., and Sachse, C. (2025). Structural basis for Vipp1 membrane binding: from loose coats and carpets to ring and rod assemblies. Nat Struct Mol Biol 32, 555–570.

Junglas, B., Orru, R., Axt, A., Siebenaller, C., Steinchen, W., Heidrich, J., Hellmich, U.A., Hellmann, N., Wolf, E., Weber, S.A.L., et al. (2020). IM30 IDPs form a membrane protective carpet upon super-complex disassembly. Commun Biol 3, 595.

Junglas, B., and Schneider, D. (2018). What is Vipp1 good for? Mol Microbiol 108, 1–5.

Kelly, C.L., Taylor, G.M., Hitchcock, A., Torres-Mendez, A., and Heap, J.T. (2018). A Rhamnose-Inducible System for Precise and Temporal Control of Gene Expression in Cyanobacteria. ACS synthetic biology 7, 1056–1066.

Kozak, M., and Kaksonen, M. (2022). Condensation of Ede1 promotes the initiation of endocytosis. eLife 11.

Kroll, D., Meierhoff, K., Bechtold, N., Kinoshita, M., Westphal, S., Vothknecht, U.C., Soll, J., and Westhoff, P. (2001). VIPP1, a nuclear gene of Arabidopsis thaliana essential for thylakoid membrane formation. Proc Natl Acad Sci U S A 98, 4238–4242.

Lasker, K., von Diezmann, L., Zhou, X., Ahrens, D.G., Mann, T.H., Moerner, W.E., and Shapiro, L. (2020). Selective sequestration of signalling proteins in a membraneless organelle reinforces the spatial regulation of asymmetry in Caulobacter crescentus. Nature microbiology 5, 418–429.

Li, H.M., Kaneko, Y., and Keegstra, K. (1994). Molecular cloning of a chloroplastic protein associated with both the envelope and thylakoid membranes. Plant Mol Biol 25, 619–632.

Linsenmeier, M., Hondele, M., Grigolato, F., Secchi, E., Weis, K., and Arosio, P. (2022). Dynamic arrest and aging of biomolecular condensates are modulated by low-complexity domains, RNA and biochemical activity. Nat Commun 13.

Liu, C., Willmund, F., Golecki, J.R., Cacace, S., Hess, B., Markert, C., and Schroda, M. (2007). The chloroplast HSP70B-CDJ2-CGE1 chaperones catalyse assembly and disassembly of VIPP1 oligomers in Chlamydomonas. Plant J 50, 265–277.

Liu, J., Tassinari, M., Souza, D.P., Naskar, S., Noel, J.K., Bohuszewicz, O., Buck, M., Williams, T.A., Baum, B., and Low, H.H. (2021). Bacterial Vipp1 and PspA are members of the ancient ESCRT-III membrane-remodeling superfamily. Cell 184, 3660–3673 e3618.

Liu, Y., Lee, Y.J., and Cicerone, M.T. (2009). Broadband CARS spectral phase retrieval using a time-domain Kramers-Kronig transform. Optics letters 34, 1363–1365.

Lyon, A.S., Peeples, W.B., and Rosen, M.K. (2021). A framework for understanding the functions of biomolecular condensates across scales. Nat Rev Mol Cell Biol 22, 215–235.

Ma, W., and Mayr, C. (2018). A Membraneless Organelle Associated with the Endoplasmic Reticulum Enables 3’UTR-Mediated Protein-Protein Interactions. Cell 175, 1492–1506 e1419.

Maity, S., Caillat, C., Miguet, N., Sulbaran, G., Effantin, G., Schoehn, G., Roos, W.H., and Weissenhorn, W. (2019). VPS4 triggers constriction and cleavage of ESCRT-III helical filaments. Science advances 5, eaau7198.

Mangiarotti, A., Chen, N.N., Zhao, Z.L., Lipowsky, R., and Dimova, R. (2023). Wetting and complex remodeling of membranes by biomolecular condensates. Nat Commun 14.

Martinez, K.A., 2nd, Kitko, R.D., Mershon, J.P., Adcox, H.E., Malek, K.A., Berkmen, M.B., and Slonczewski, J.L. (2012). Cytoplasmic pH response to acid stress in individual cells of Escherichia coli and Bacillus subtilis observed by fluorescence ratio imaging microscopy. Appl Environ Microbiol 78, 3706-3714.

Mathieu, C., Pappu, R.V., and Taylor, J.P. (2020). Beyond aggregation: Pathological phase transitions in neurodegenerative disease. Science 370, 56–60.

McCullough, J., Frost, A., and Sundquist, W.I. (2018). Structures, Functions, and Dynamics of ESCRT-III/Vps4 Membrane Remodeling and Fission Complexes. Annu Rev Cell Dev Biol 34, 85-109.

Mittag, T., and Pappu, R.V. (2022). A conceptual framework for understanding phase separation and addressing open questions and challenges. Mol Cell 82, 2201–2214.

Mullineaux, C.W. (2014). Co-existence of photosynthetic and respiratory activities in cyanobacterial thylakoid membranes. Biochim Biophys Acta 1837, 503–511.

Murthy, A.C., Dignon, G.L., Kan, Y., Zerze, G.H., Parekh, S.H., Mittal, J., and Fawzi, N.L. (2019). Molecular interactions underlying liquid-liquid phase separation of the FUS low-complexity domain. Nat Struct Mol Biol 26, 637–648.

Muthunayake, N.S., Tomares, D.T., Childers, W.S., and Schrader, J.M. (2020). Phase-separated bacterial ribonucleoprotein bodies organize mRNA decay. Wiley interdisciplinary reviews RNA 11, e1599.

Naskar, S., Merino, A., Espadas, J., Singh, J., Roux, A., Colom, A., and Low, H.H. (2025). Mechanism for Vipp1 spiral formation, ring biogenesis, and membrane repair. Nat Struct Mol Biol 32, 571–584.

Nguyen, H.C., Talledge, N., McCullough, J., Sharma, A., Moss, F.R., 3rd, Iwasa, J.H., Vershinin, M.D., Sundquist, W.I., and Frost, A. (2020). Membrane constriction and thinning by sequential ESCRT-III polymerization. Nat Struct Mol Biol 27, 392-399.

O’Flynn, B.G., and Mittag, T. (2021). The role of liquid-liquid phase separation in regulating enzyme activity. Curr Opin Cell Biol 69, 70–79.

Obita, T., Saksena, S., Ghazi-Tabatabai, S., Gill, D.J., Perisic, O., Emr, S.D., and Williams, R.L. (2007). Structural basis for selective recognition of ESCRT-III by the AAA ATPase Vps4. Nature 449, 735–739.

Padan, E., Bibi, E., Ito, M., and Krulwich, T.A. (2005). Alkaline pH homeostasis in bacteria: New insights. Bba-Biomembranes 1717, 67–88.

Pan, S., Gries, K., Engel, B.D., Schroda, M., Haselwandter, C.A., and Scheuring, S. (2024). The cyanobacterial protein VIPP1 forms ESCRT-III-like structures on lipid bilayers. Nat Struct Mol Biol.

Pappu, R.V., Cohen, S.R., Dar, F., Farag, M., and Kar, M. (2023). Phase Transitions of Associative Biomacromolecules. Chem Rev 123, 8945–8987.

Pfitzner, A.K., Moser von Filseck, J., and Roux, A. (2021). Principles of membrane remodeling by dynamic ESCRT-III polymers. Trends Cell Biol 31, 856–868.

Portz, B., and Shorter, J. (2018). 3’ UTRs in the Eye of the TIGER. Dev Cell 47, 544–546.

Quarta, N., Bhandari, T.R., Debrich, K., Hellmann, N., Girard, M., and Schneider, D. (2026). The cyanobacterial ESCRT-III protein IM30 forms biomolecular condensates at physiologically relevant conditions. Biophys J 125, 1081–1094.

Quarta, N., Bhandari, T.R., Girard, M., Hellmann, N., and Schneider, D. (2024). Monomer unfolding of a bacterial ESCRT-III superfamily member is coupled to oligomer disassembly. Protein Sci 33, e5187.

Racki, L.R., Tocheva, E.I., Dieterle, M.G., Sullivan, M.C., Jensen, G.J., and Newman, D.K. (2017). Polyphosphate granule biogenesis is temporally and functionally tied to cell cycle exit during starvation in Pseudomonas aeruginosa. Proc Natl Acad Sci U S A 114, E2440–E2449.

Ramasubramani, V., Adorf, C.S., Dodd, P.M., Dice, B.D., and Glotzer, S.C. (2018). signac: A Python framework for data and workflow management. Paper presented at: 17th Python in Science Conference.

Ramirez, D.A., Hough, L.E., and Shirts, M.R. (2024). Coiled-coil domains are sufficient to drive liquid-liquid phase separation in protein models. Biophys J 123, 703–717.

Ray, S., Singh, N., Kumar, R., Patel, K., Pandey, S., Datta, D., Mahato, J., Panigrahi, R., Navalkar, A., Mehra, S., et al. (2020). alpha-Synuclein aggregation nucleates through liquid-liquid phase separation. Nature chemistry 12, 705–716.

Remec Pavlin, M., and Hurley, J.H. (2020). The ESCRTs - converging on mechanism. J Cell Sci 133.

Rippka, R., Deruelles, J., Waterbury, J.B., Herdman, M., and Stanier, R.Y. (1979). Generic assignments, strains histories and properties of pure cultures of cyanobacteria. J Gen Microbiol 111, 1–61.

Rubinstein, M., and Colby, R.H. (2003). Polymer Physics (Oxford University Press).

Rueden, C.T., Schindelin, J., Hiner, M.C., DeZonia, B.E., Walter, A.E., Arena, E.T., and Eliceiri, K.W. (2017). ImageJ2: ImageJ for the next generation of scientific image data. BMC bioinformatics 18, 529.

Sasazawa, M., Tomares, D.T., Childers, W.S., and Saurabh, S. (2024). Biomolecular condensates as stress sensors and modulators of bacterial signaling. PLoS pathogens 20, e1012413.

Saur, M., Hennig, R., Young, P., Rusitzka, K., Hellmann, N., Heidrich, J., Morgner, N., Markl, J., and Schneider, D. (2017). A Janus-Faced IM30 Ring Involved in Thylakoid Membrane Fusion Is Assembled from IM30 Tetramers. Structure 25, 1380–1390 e1385.

Schindelin, J., Arganda-Carreras, I., Frise, E., Kaynig, V., Longair, M., Pietzsch, T., Preibisch, S., Rueden, C., Saalfeld, S., Schmid, B., et al. (2012). Fiji: an open-source platform for biological-image analysis. Nature methods 9, 676-682.

Schlösser, L., Kutzner, M., Hellmann, N., Kiesewetter, D., Bieber, J., Quarta, N., Ge, X., Goetze, T., Junglas, B., Matsumura, F., et al. (2025). Membrane binding of a cyanobacterial ESCRT-III protein crucially involves the helix alpha1-3 hairpin conserved in all superfamily members. Protein Sci 34, e70387.

Schlosser, L., Sachse, C., Low, H.H., and Schneider, D. (2023). Conserved structures of ESCRT-III superfamily members across domains of life. Trends Biochem Sci 48, 993–1004.

Schlösser, L., Sachse, C., Low, H.H., and Schneider, D. (2023). Conserved structures of ESCRT-III superfamily members across domains of life. Trends Biochem Sci 48, 993–1004.

Shim, S., Kimpler, L.A., and Hanson, P.I. (2007). Structure/function analysis of four core ESCRT-III proteins reveals common regulatory role for extreme C-terminal domain. Traffic 8, 1068–1079.

Shin, Y., and Brangwynne, C.P. (2017). Liquid phase condensation in cell physiology and disease. Science 357.

Siebenaller, C., Junglas, B., Lehmann, A., Hellmann, N., and Schneider, D. (2020). Proton Leakage Is Sensed by IM30 and Activates IM30-Triggered Membrane Fusion. Int J Mol Sci 21.

Siebenaller, C., Junglas, B., and Schneider, D. (2019). Functional Implications of Multiple IM30 Oligomeric States. Front Plant Sci 10, 1500.

Siebenaller, C., Schlosser, L., Junglas, B., Schmidt-Dengler, M., Jacob, D., Hellmann, N., Sachse, C., Helm, M., and Schneider, D. (2021). Binding and/or hydrolysis of purine-based nucleotides is not required for IM30 ring formation. FEBS Lett 595, 1876–1885.

Sing, C.E., and Perry, S.L. (2020). Recent progress in the science of complex coacervation. Soft matter 16, 2885–2914.

Slonczewski, J.L., Fujisawa, M., Dopson, M., and Krulwich, T.A. (2009). Cytoplasmic pH Measurement and Homeostasis in Bacteria and Archaea. Adv Microb Physiol 55, 1–79.

Soumpasis, D.M. (1983). Theoretical analysis of fluorescence photobleaching recovery experiments. Biophys J 41, 95–97.

Staples, M.I., Frazer, C., Fawzi, N.L., and Bennett, R.J. (2023). Phase separation in fungi. Nature microbiology 8, 375–386.

Stevens, D.A., and Lasker, K. (2025). Microbial biomolecular condensates: from conserved principles to synthetic biology opportunities. Curr Opin Microbiol 88, 102677.

Su, Q., Mehta, S., and Zhang, J. (2021). Liquid-liquid phase separation: Orchestrating cell signaling through time and space. Molecular Cell 81, 4137–4146.

Tan, J., Davies, B.A., Payne, J.A., Benson, L.M., and Katzmann, D.J. (2015). Conformational Changes in the Endosomal Sorting Complex Required for the Transport III Subunit Ist1 Lead to Distinct Modes of ATPase Vps4 Regulation. J Biol Chem 290, 30053–30065.

Taylor, N.O., Wei, M.T., Stone, H.A., and Brangwynne, C.P. (2019). Quantifying Dynamics in Phase-Separated Condensates Using Fluorescence Recovery after Photobleaching. Biophys J 117, 1285–1300.

Tesei, G., and Lindorff-Larsen, K. (2022). Improved predictions of phase behaviour of intrinsically disordered proteins by tuning the interaction range. Open research Europe 2, 94.

Tesei, G., Trolle, A.I., Jonsson, N., Betz, J., Knudsen, F.E., Pesce, F., Johansson, K.E., and Lindorff-Larsen, K. (2024). Conformational ensembles of the human intrinsically disordered proteome. Nature 626.

Thurotte, A., and Schneider, D. (2019). The Fusion Activity of IM30 Rings Involves Controlled Unmasking of the Fusogenic Core. Front Plant Sci 10, 108.

Van Lindt, J., Bratek-Skicki, A., Nguyen, P.N., Pakravan, D., Duran-Armenta, L.F., Tantos, A., Pancsa, R., Van Den Bosch, L., Maes, D., and Tompa, P. (2021). A generic approach to study the kinetics of liquid-liquid phase separation under near-native conditions. Commun Biol 4, 77.

Vietri, M., Radulovic, M., and Stenmark, H. (2020). The many functions of ESCRTs. Nat Rev Mol Cell Biol 21, 25–42.

Wang, H., Yan, X., Aigner, H., Bracher, A., Nguyen, N.D., Hee, W.Y., Long, B.M., Price, G.D., Hartl, F.U., and Hayer-Hartl, M. (2019). Rubisco condensate formation by CcmM in beta-carboxysome biogenesis. Nature 566, 131–135.

Westphal, S., Heins, L., Soll, J., and Vothknecht, U.C. (2001). Vipp1 deletion mutant of Synechocystis: a connection between bacterial phage shock and thylakoid biogenesis? Proc Natl Acad Sci USA 98, 4243–4248.

Yamaguchi, S., Reid, D.A., Rothenberg, E., and Darwin, A.J. (2013). Changes in Psp protein binding partners, localization and behaviour upon activation of the Yersinia enterocolitica phage shock protein response. Molecular Microbiology 87, 656–671.

Yamashita, T., and Voth, G.A. (2010). Properties of hydrated excess protons near phospholipid bilayers. J Phys Chem B 114, 592–603.

Zeng, M., Shang, Y., Araki, Y., Guo, T., Huganir, R.L., and Zhang, M. (2016). Phase Transition in Postsynaptic Densities Underlies Formation of Synaptic Complexes and Synaptic Plasticity. Cell 166, 1163–1175 e1112.

Zeng, M.L., Chen, X.D., Guan, D.S., Xu, J., Wu, H.W., Tong, P.E., and Zhang, M.J. (2018). Reconstituted Postsynaptic Density as a Molecular Platform for Understanding Synapse Formation and Plasticity. Cell 174, 1172-+.

Zhang, L., Kato, Y., Otters, S., Vothknecht, U.C., and Sakamoto, W. (2012). Essential Role of VIPP1 in Chloroplast Envelope Maintenance in Arabidopsis. Plant Cell 24, 3695–3707.

Zhou, H.B., Huertas, J., Maristany, M.J., Russell, K., Hwang, J.H., Yao, R.W., Samanta, N., Hutchings, J., Billur, R., Shiozaki, M., et al. (2025). Multiscale structure of chromatin condensates explains phase separation and material properties. Science 390.

