## Supplemental Figures S1-S8 for "The structured hairpin region of the bacterial ESCRT-III protein IM30 orchestrates stress-induced condensate formation"

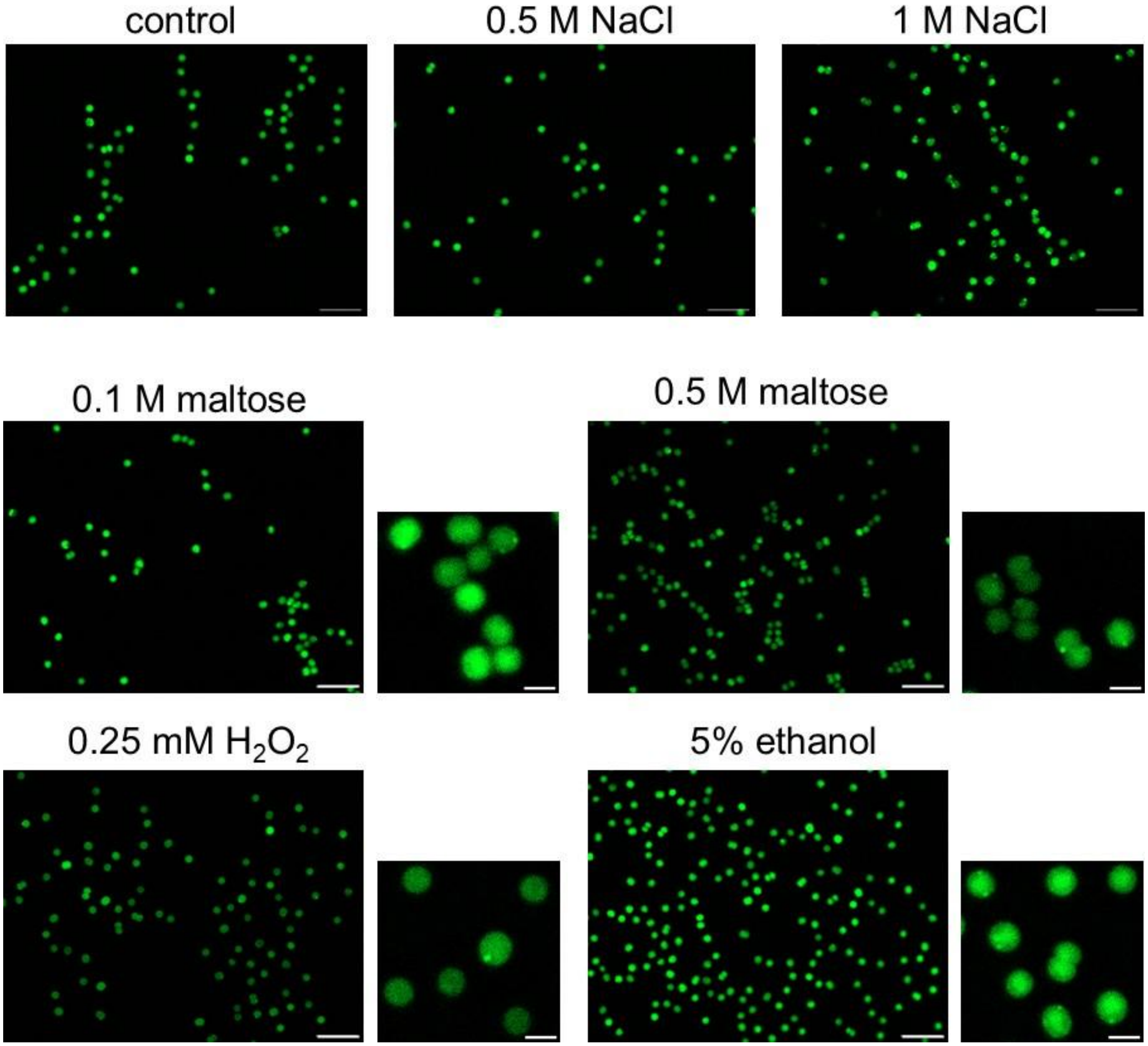
control
0.5 M NaCl
1 M NaCl
0.1 M maltose
0.5 M maltose
0.25 mM $H_2O_2$
5% ethanol

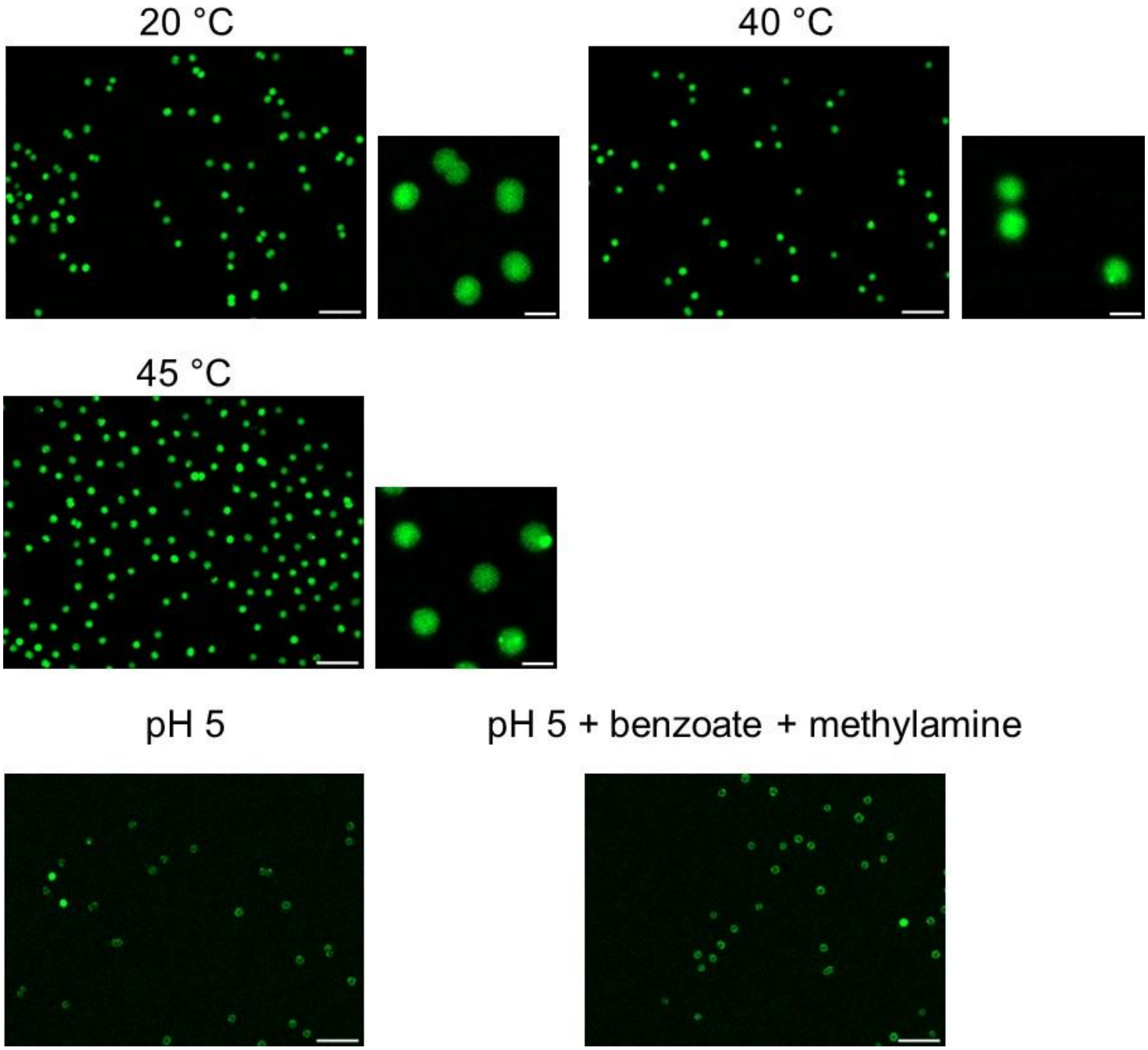


**Figure S1.** Representative overview images of stressed *Synechocystis* cells.

Stress conditions are described in the main text. Magnified views are omitted here where already presented in the main figures.

Scale bars: 10 µm (overview images); 2 µm (zoomed panels).

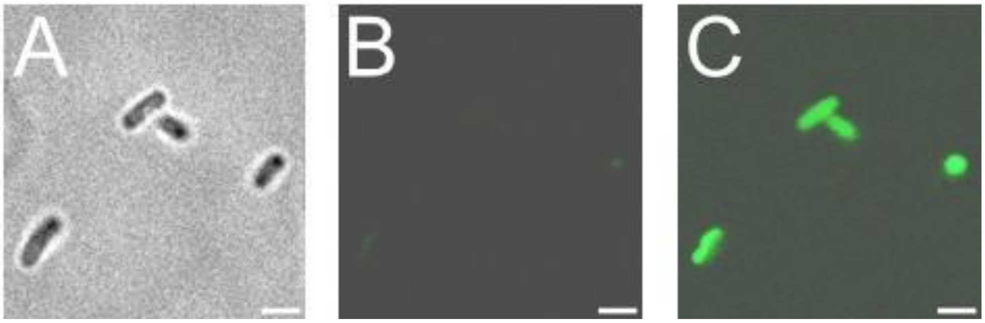


**Figure S2.** mVenus signal in non-induced *E. coli* cells harboring the plasmid for wt IM30-mVenus expression.

(A) Phase contrast image. (B, C) Under standard imaging conditions, no significant background fluorescence was detectable prior to IPTG induction (B; cf. Figure 2B in the main text). However, upon increasing image contrast, a uniform, low-level mVenus signal was discernible throughout the cytoplasm of all cells, even without induction (C).

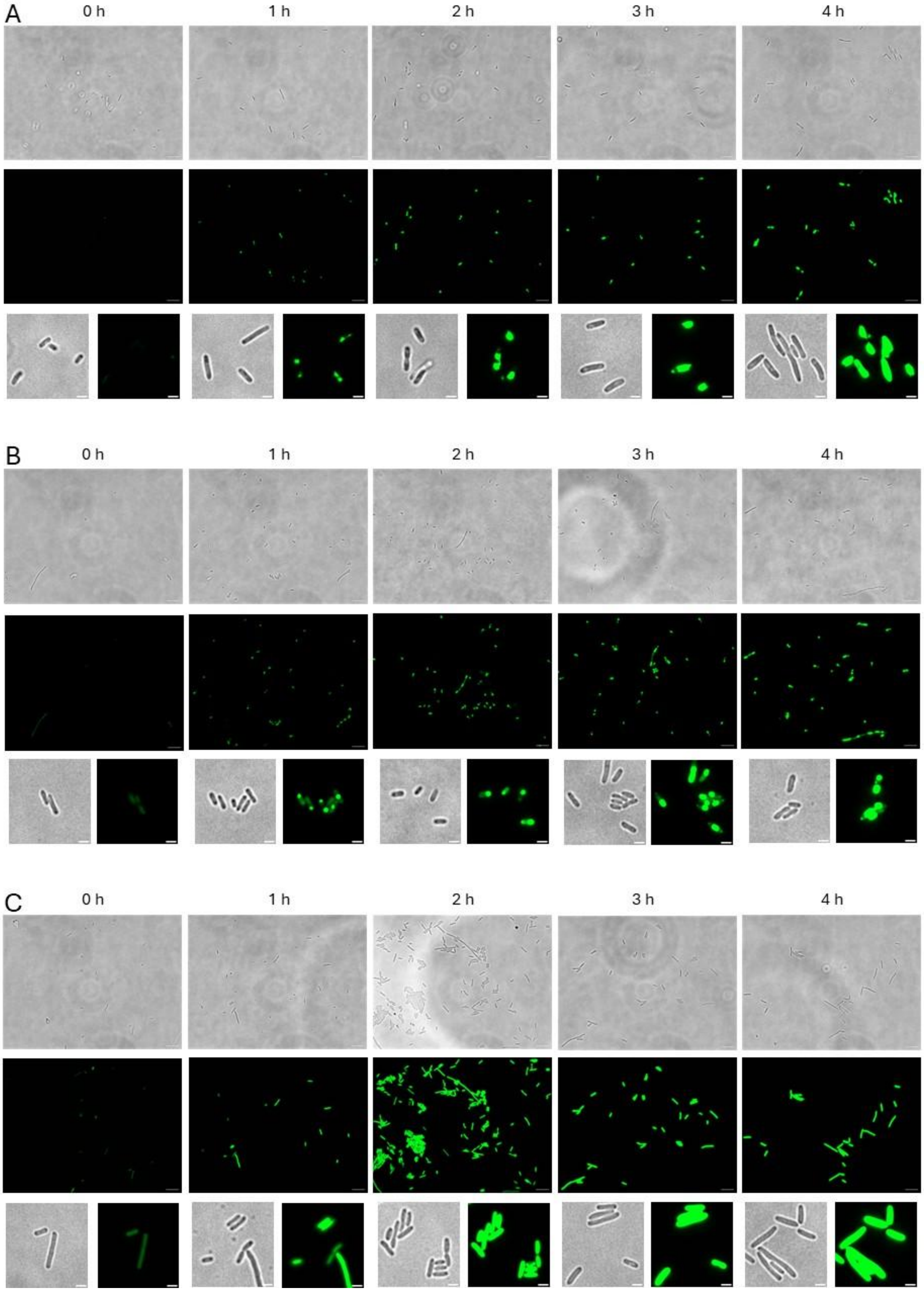
A
0 h
1 h
2 h
3 h
4 h
B
0 h
1 h
2 h
3 h
4 h
C
0 h
1 h
2 h
3 h
4 h

**Figure S3:** Non-merged overview images showing multiple *E. coli* cells.

*E. coli* cells expressing (A) wt IM30-mVenus, (B) IM30*-mVenus, or (C) free mVenus were imaged prior to induction (0 h) and at hourly intervals following the addition of 0.5 mM IPTG to induce gene expression. Scale bars: 10 µm (overview images); 2 µm (zoomed images).

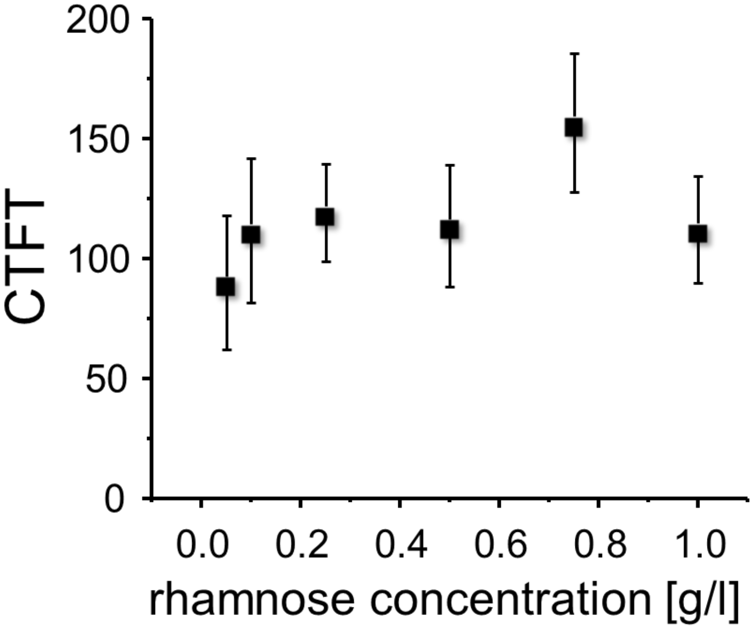


**Figure S4.** Corrected Total Cell Fluorescence (CTCF) of IM30-mVenus–expressing cyanobacteria under increasing concentrations of L-rhamnose.

CTCF was calculated for individual cells using ImageJ. For each cell, the area and integrated density were measured and corrected for background fluorescence according to:

CTCF = Integrated Density – Area × Mean Background Fluorescence.

At least 10 cells per condition were analyzed to ensure statistical robustness.

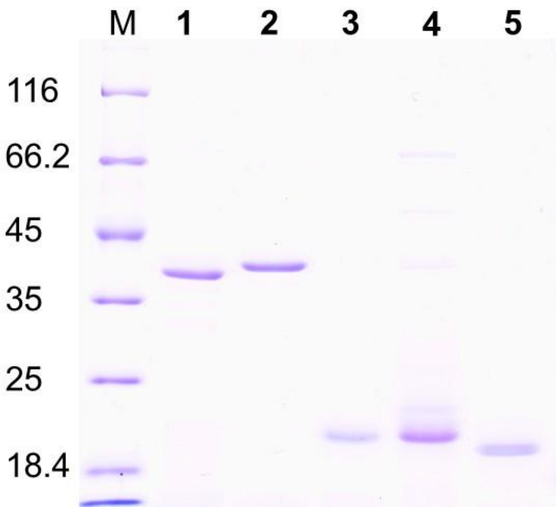


**Figure S5.** SDS-PAGE analysis of purified proteins used in this study.

Lanes: M, molecular weight marker (masses indicated on the left); 1: wt IM30; 2: IM30*: 3, IM30 α0–3; 4: IM30 α4–6; 5: IM30 α1–3.

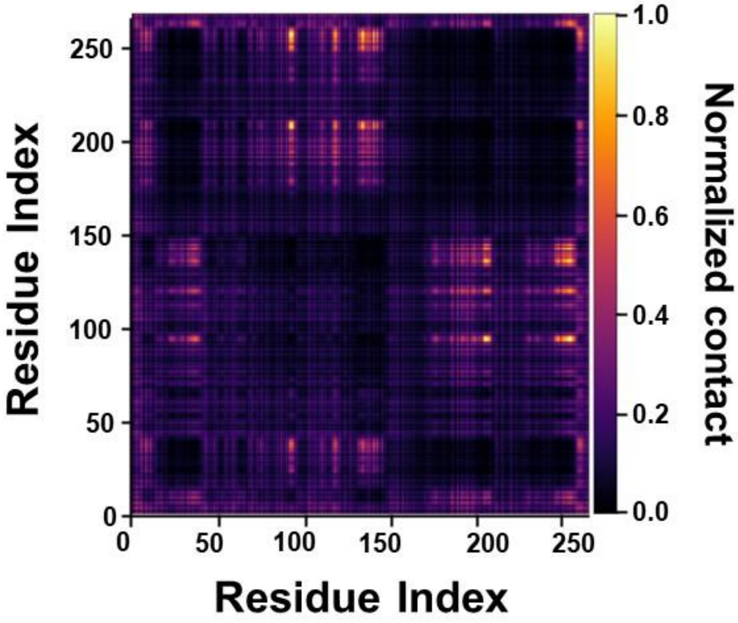


**Figure S6**: Interchain contact map of IM30 condensates at 220 K and $\kappa = 0.097$ Å$^{-1}$ for the full-length IM30*.

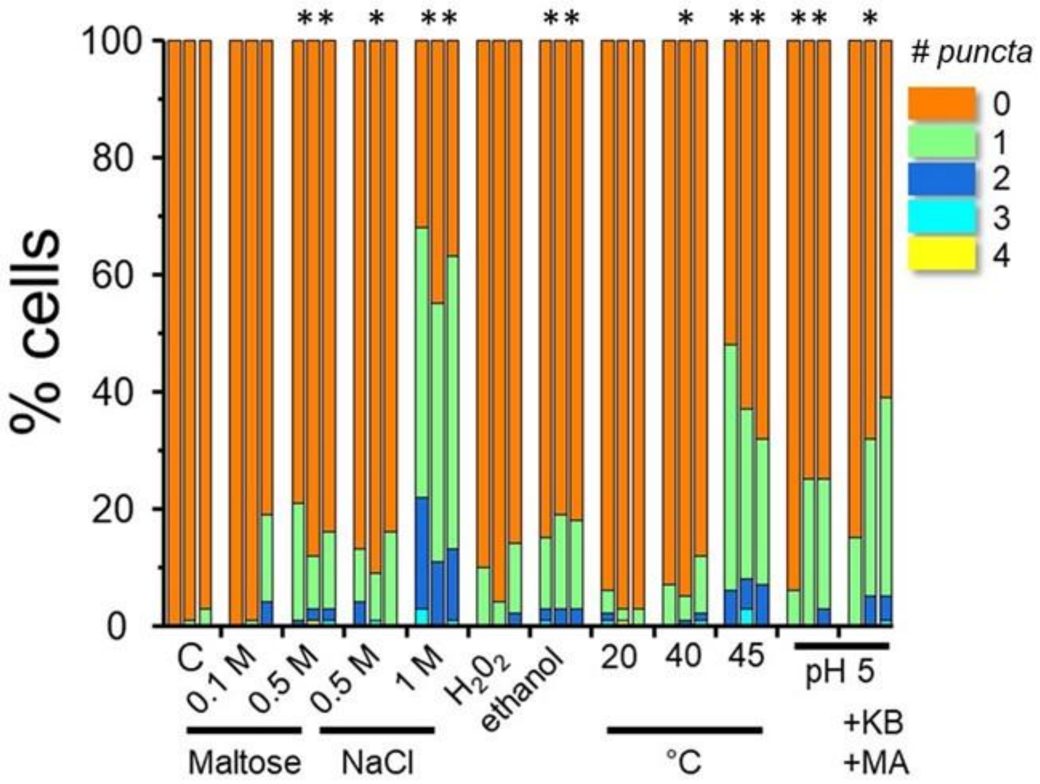


**Figure S7**: Stress-induced formation of IM30 *puncta* in *Synechocystis*.

*Synechocystis* cells expressing IM30–mVenus were subjected to the indicated stress conditions (see Methods for details). Fifteen minutes after stress application, cells were imaged and analyzed. The proportion of cells exhibiting *puncta* was quantified. Here, the individual results of three biological replicates are shown. KB: potassium benzoate; MA: methylamine hydrochloride. Statistical significance was assessed by two-tailed unpaired t-test relative to unstressed control: $^{**}p \leq 0.001$, $^{*}p \leq 0.05$.

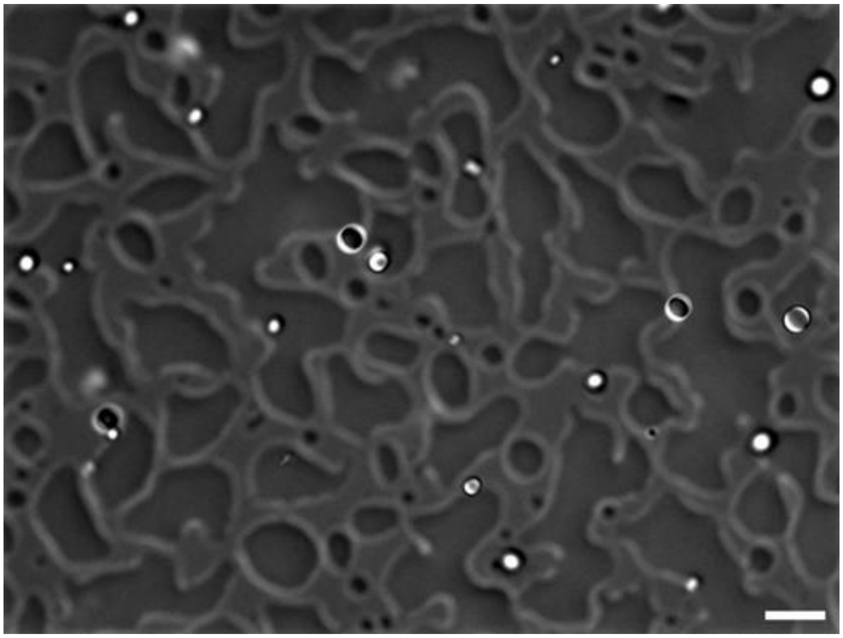

**Figure S8**: Wetting of pH-induced IM30* condensates on glass surfaces.

An exemplary DIC image showing wetting of bottom glass surface by IM30* condensates after prolonged incubation time. Condensates were formed at pH 5.5, in 10 mM HEPES and 10 mM Pi-buffer, in the absence of NaCl or PEG. Scale bar: 10 µm.
